# A new (old) approach to genotype-based phylogenomic inference within species, with an example from the saguaro cactus (*Carnegiea gigantea*)

**DOI:** 10.1101/2020.06.17.157768

**Authors:** Michael J. Sanderson, Alberto Búrquez, Dario Copetti, Michelle M. McMahon, Yichao Zeng, Martin F. Wojciechowski

## Abstract

Genome sequence data are routinely being used to infer phylogenetic history within and between closely related diploid species, but few tree inference methods are specifically tailored to diploid genotype data. Here we re-examine the method of “polymorphism parsimony” (Inger 1967; Farris 1978; Felsenstein 1979), originally introduced to study morphological characters and chromosome inversion polymorphisms, to evaluate its utility for unphased diploid genotype data in large scale phylogenomic data sets. We show that it is equivalent to inferring species trees by minimizing deep coalescences—assuming an infinite sites model. Two potential advantages of this approach are scalability and estimation of a rooted tree. As with some other single nucleotide polymorphism (SNP) based methods, it requires thinning of data sets to statistically independent sites, and we describe a genotype-based test for phylogenetic independence. To evaluate this approach in genome scale data, we construct intraspecific phylogenies for 10 populations of the saguaro cactus using 200 Gbp of resequencing data, and then use these methods to test whether the population with highest genetic diversity corresponds to the root of the genotype trees. Results were highly congruent with the (unrooted) trees obtained using SVDquartets, a scalable alternative method of phylogenomic inference.

---

Most published phylogenetic trees of eukaryotes have diploid taxa at their leaves, but most phylogenetic methods implicitly assume input sequence data are haploid—a single row of individual character states, such as nucleotide base calls, for each leaf label. The significance of this disconnect is growing, especially in studies that include multiple individuals within species *and* have high enough coverage to call diploid genotypes reliably. Judging by the number of recent reconstructions of phylogenies of individual diploid accessions within species, there is a gowing demand to characterize the phylogenetic structure in these genotype data (e.g. Durvasula et al. 2017; Wang et al. 2018; Zhao et al. 2019). Yet, even recent comprehensive reviews on phylogenomics (Bravo et al. 2019; Liu et al. 2019) make little mention of “genotypes”, despite the fact that diploidy, and polyploidy in general, allows intra-individual polymorphism, which is a well known complicating factor in phylogenetic inference long known in gene families, for example (Goodman et al. 1979; Potts et al. 2014).

Various “workarounds” have been used to recode genotype data into a form that can be used by existing phylogenetic methods. In maximum parsimony, for example, the three genotypes for a biallelic SNP, {*AA, Aa, aa*}, can be recoded (i) as the integer states {0, 1, 2} and treat them as either unordered or ordered multistate characters (Rheindt et al. 2014), possibly with weighting (Buckler and Holtsford 1996); (ii) as the states {0, ?, 1}, treating the heterozygote as missing; (iii) as the states {0, {0, 1}, 1}, where the heterozygote is treated as “polymorphic” (in the sense of PAUP* or MacClade, which does not allow an ancestor to be polymorphic, Maddison and Maddison 2000); or (iv) as a pair of binary presence-absence characters, one per allele (Schmidt-Lebuhn et al. 2017). Some of these recoding schemes can be used in likelihood inference (e.g., method (i) in VCFtoTree: Xu et al. 2017) or distance methods. The widely used identity-by-state (IBS) distance (Chang et al. 2015; Subramanian et al. 2019) is actually the same as that implied by the ordered integer coding in (i) above when applied to a single site. It is also the same as the Manhattan distance in allele frequency space if the “population” is just the two alleles in a diploid individual.The latter was used for individual site branch lengths in the FreqPars method of Swofford and Berlocher (1987).

Three other frameworks are more explicit about the nature of evolving genotypes. The method of *polymorphism parsimony* (PP) (Inger 1967; Farris 1978; Felsenstein 1979, 2004, 2005) infers phylogenies from discrete characters that can be ancestrally polymorphic by minimizing the extent of polymorphism on the tree. Though originally applied to morphological traits and chromosome inversions, it seems clearly applicable to diploid biallelic genotypes derived from genome sequence data. The complexity of morphological traits and chromosome structure led these authors to invoke specific assumptions about character evolution: a unique derivation with no losses of the derived state. This allowed polymorphism to be gained once but lost many times. These assumptions turn out to mirror the infinite sites model of sequence evolution (Kimura 1969), a similarity we return to below. PP is implemented in Phylip (Felsenstein 2005) but has has been used infrequently (Baum 1983; Suh et al. 2015).

Two other frameworks, gene tree reconciliation in phylogenetics (Goodman et al. 1979; Page 1994), and coalescent theory in population genetics (Kingman 1982; Hein et al. 2005), also provide toolkits for modeling genotypes more explicitly by analyzing the genealogy of alleles. Yet, existing implementations are rather agnostic about using genotype vs. haplotype data. For example, considerable headway has been made in tying phylogenetic inference between species/populations to allele phylogenies using either the gene tree reconciliation framework, which aims to infer species/population trees by minimizing the deep coalescence (DC) score in allele trees (Maddison 1997; Than and Nakhleh 2009); or a more explicit probabilistic model provided by the multi-species coalescent (MSC) (Degnan and Rosenberg 2009). In the MSC framework, several methods use biallelic SNP data to build trees, such as SNAPP (Bryant et al. 2012); SVDquartets (Chifman and Kubatko 2014; Vachaspati and Warnow 2018); Phylonet (Zhu and Nakhleh 2018). Yet, none of these methods embrace genotype data per se. Instead, they assign single sequences—haplotypes—to predefined populations or species (possibly one sequence per species) and infer the phylogenies of those taxa.

These approaches (and others) can be “spoofed” into taking diploid genotype inputs by converting them to pairs of pseudo-haplotypes and then aggregating them into populations or species. At the lowest levels these would be “populations” of two pseudo-haplotypes. We say “pseudo-haplotypes” unless they are explicitly phased either computationally (e.g. Chifman and Kubatko 2014) or experimentally. For example, VCFtoTree (Xu et al. 2017) can use phased haplotypes, but if the data are unphased it simply ignores heterozygous sites. Alternatively, if all sites are treated as coalescent independent sites (Tian and Kubatko 2017), then the two haplotypes in each diploid can be constructed by segregating the alternative alleles in all heterozygotes randomly.

However, this raises another issue. At the species level and below, even a phylogenetic inference method properly focused on genotypes still runs into the problem of gene flow and non-tree-like history. In a Wright-Fisher population of genotypes, the paired alleles within a diploid individual most likely do not coalesce with each other more recently than they do with alleles from other individuals in that population. In other words, individuals (genotypes) do not act like populations. However, this does not invalidate the concept of genotype tree or genotype-specific tree inference methods, per se; it means they run the same risks as many other phylogenetic tools applied to the infraspecific level.

Existing approaches to building trees with genotype data, whether workarounds, or implicit genotype-based tools, are all ad hoc to various degrees, and it may prove useful to examine all of them in a common allele tree framework to expose assumptions and limitations. In this paper we take a step toward this goal by showing conditions under which polymorphism parsimony, and allele tree reconciliation by minimizing deep coalescences, are equivalent. We then exploit the existing implementation of PP in Phylip (Felsenstein 2005) to study phylogenetic structure among genotypes of saguaro cactus (*Carnegiea gigantea*), across its range, illustrating its scalability to millions of SNPs and utility in estimating a rooted genotype tree. By adopting the allele tree framework shared by reconciliation methods and coalescent theory, we hope to illuminate the strengths of genotype data while exploring the limits to infraspecific phylogenetic inference imposed by interbreeding.

## Materials and Methods I. Theory

### Genotypes, characters and trees

Assuming each site is diploid and biallelic with alternate alleles labelled 0 and 1, and has therefore three possible diploid genotype states plus “missing”, we will use one of two notation schemes for the genotypes of individual *i* and site *j*:

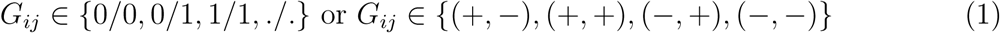

The first is the notation typical for biallelic SNPs in modern genotyping pipelines (e.g., VCF format). The second (useful in proofs in the Appendix) is a vector in which the first and second components denote the presence (+) or absence (−) of the 0 and 1 alleles, respectively, in that genotype. Define a *genotype character* for site *j* in *n* individuals as the vector, (*G*_1*j*_, …, *G*_*nj*_). We will often use the shorthand, *G*, for a genotype character.

We assume that genotype characters are statistically independent—that is, sites are *coalescent independent* (Tian and Kubatko 2017)—but we test for dependence to obtain such data. Genotypes at different sites in an individual are assumed to be *unphased* (genotypes are *phased* if the alternate alleles in heterozygous genotypes at different sites are ordered with respect to each other as *haplotypes*).

A *genotype tree*, Ψ for *n* individuals is a rooted tree with *n* leaves. A genotype character, *G*, for these individuals has an *allele tree, t*(*G*), with at most two leaves per individual (fewer if genotype data are missing), which is imbedded within Ψ, in keeping with the gene tree reconciliation framework (Goodman et al. 1979; Page 1994; Page and Charleston 1997). Both trees are assumed to be binary.

### Equivalence of Polymorphism Parsimony and Minimizing Deep Coalescence in Allele Trees

Our main theoretical result is that for any Ψ and *G*, the PP score is the same as the deep coalescence score *for an appropriately specified allele tree, t*(*G*). This underlying allele tree is always implicitly present, but its coalescent history (Degnan and Salter 2005) and therefore deep coalescence score, are not identifiable without additional constraints, which we discuss below.

### Polymorphism parsimony for genotype

Polymorphism parsimony assumes a particular model underling the evolution of genotypes at a site: (i) the 0/0 genotype is ancestral; (ii) the 0/1 genotype can originate only once, and (iii) both 0/0 and 1/1 can evolve from 0/1 any number of times (Farris 1978; Felsenstein 1979). The original rationale was that a novel state, such as a new chromosome inversion, arises once in an ancestral edge of the tree, at which point the edge is polymorphic along with the ancestral chromosome type, and then later the *polymorphism* may be repeatedly lost, by fixation of one or the other type, but, importantly, the novel trait itself is never reversed.

Let 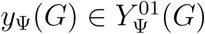 be a set of ancestral states that jointly satisfy these assumptions for genotype character, *G*, where 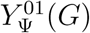 is the set of all such joint reconstructions. The number of polymorphic edges, *m*(*y*_Ψ_(*G*)), which is the number of edges having 0/1 genotypes at both ends, provides an optimality criterion.

The *polymorphism parsimony (PP) score* is then found by solving the following problem:

### Problem (PP Score)

Given Ψ and *G*, find the ancestral state set, *y*_*opt*_ over all 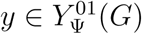 that minimizes *m*(*y*). Then the PP score is

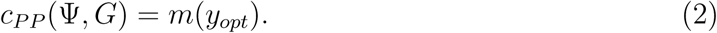

Felsenstein (1979) outlined a two-pass algorithm to compute these ancestral states and the PP score. Details are provided in the Appendix, but the intuition is as follows: in the downpass each node is assigned a pair of presence/absence flags that indicate whether allele 0 and 1, respectively, are found among any descendant leaves. Because 0 is ancestral, this immediately means that the 0 allele is present at this node. However, the 1 allele might not yet have evolved. The second phase is an uppass from the root, delaying as long as possible the final assignment of the first appearance of a 1 allele. This ensures a minimal number of edges of the tree having both 0 and 1. Figure 1 illustrates an example.

**Fig. 1.**
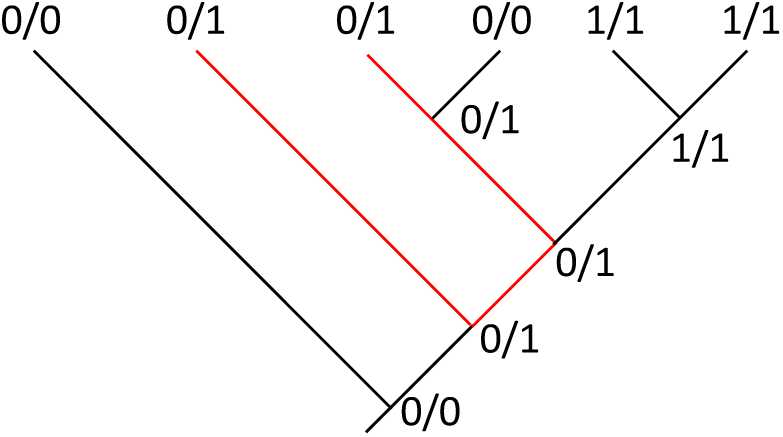
Ancestral state reconstruction under polymorphism parsimony. The ancestral states are jointly valid under the assumptions of the PP model. Optimal reconstruction indicates four polymorphic edges (red lines) and therefore a PP score of four.

PP can return different ancestral state reconstructions depending on whether 0/0 or 1/1 is assumed to be the ancestral genotype. The Phylip implementation computes the PP score for both choices and selects the smaller of the two. Although neither Farris nor Felsenstein explicitly consider missing data in their papers, the modification to the Felsenstein algorithm is slight (see Appendix). The Felsenstein algorithm runs in *O*(*n*) time, where *n* is the number of leaves of Ψ.

### Allele Tree Framework

Minimizing the number of polymorphic edges in a tree of genotypes sounds suspiciously like “minimizing deep coalescences”, but the latter usually refers to gene trees imbedded within species or population trees. The connection between these seemingly different frameworks can be made quite strong for genotype data under suitable assumptions. First, we assume the existence of a genotype tree, Ψ, containing an allele tree, *t*(*G*), giving rise to the observed genotypes at the leaves of Ψ. Second, because any allele tree can be concordant with any genotype tree trivially by allowing some or all coalescences to occur below the root (Degnan and Salter 2005, p. 25), we will constrain the allele tree to evolve according to the infinite sites model in population genetics. This assumption is consistent with the genotype evolution model described above for polymorphism parsimony. Finally, we will invoke an optimality criterion to select among all allele trees that fit this model.

The first assumption is operational. Even in a randomly mating population of individuals, we can postulate the existence of Ψ, but it would bear little relationship to the allele tree, because coalescence events would occur at random, and could therefore be imbedded in Ψ only by forcing most coalescences to occur below the root of Ψ. However, with increasing amounts of historical isolation between these individuals, there will be a bias in the distribution of allele trees imbedded within the genotype tree away from expectations of a random coalescent.

The second assumption requires allele trees, *t*, that obey the PP model. Because the three model assumptions stated above for PP now involve an interplay between *t* and Ψ in this allele tree framework, it is helpful to leverage the gene tree reconciliation framework from here on. The key model assumption is that allele states 0 and 1 on *t* are constrained such that there can only be one 0 → 1 transition and no 1 → 0 transitions. A second model assumption is that this unique origin of the 1 allele arises just “after” a coalescence event in the allele tree (i.e., more recently). This corresponds to the PP assumption that when the novel allele arises it is in a polymorphic condition in Ψ—that is, there is co-occurring allele tree edge within Ψ having the 0 allele. In the reconciliation framework this can be thought of as a “duplication”. Finally, from a polymorphic edge of Ψ, in which allele tree edges having both the 0 and 1 alleles are imbedded, one or the other allele tree edge can go extinct, or be “lost”, leaving the remaining allele tree edge present. Formally, these duplication and loss events are the same as those described in gene tree reconciliation (Zhang 2011). They are also equivalent to the infinite sites model in population genetics (Kimura 1969). Let 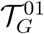 be the set of all rooted binary allele trees that (i) have leaf states with the correct mapping from *G* to binary allele states; and (ii) satisfy the infinite sites assumption.

Finally, as an optimality criterion to select among allele trees 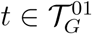, we use the *deep coalescence* (DC) score. An allele tree’s DC score, *c*_DC_(Ψ, *t*), is the total number of “extra” allele tree edges per edge of Ψ (Maddison 1997). See Appendix for further details. Now we are in a position to generalize the DC score to genotypes by solving the following:

### Problem (DC-G Score)

Given Ψ and *G*, find the allele tree, *t*_*min*_ over all 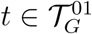 that minimizes *c*_DC_(Ψ, *t*). The corresponding DC-G (deep coalescence–genotype) score is

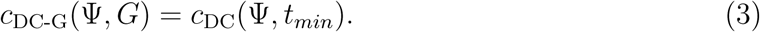

The solution, *t*_*min*_, can be found efficiently because of the constraints of the genotype evolution model (Fig. 2). Intuitively, because the 1 allele evolves exactly once and is unreversed, *t*_*min*_ has to have two subtrees, 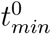 and 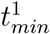, each having only leaves labelled with one of the alleles. In addition, 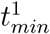 is a clade. The subtree 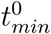 must be present at the root of Ψ, because 0 is assumed ancestral and must be present there. Moreover, to minimize the number of edges of Ψ in which both allele subtrees are present (and hence have an “extra edge” in the sense counted by the DC score) all nodes in *t*_*min*_ are placed as close to the leaves as possible. Finally, a homozygous genotype state at a leaf is represented by a coalescence in the allele tree at the leaf node, which ensures that no deep coalescence occurs on a terminal edge having a homozygous leaf (see Appendix for details).

**Fig. 2.**
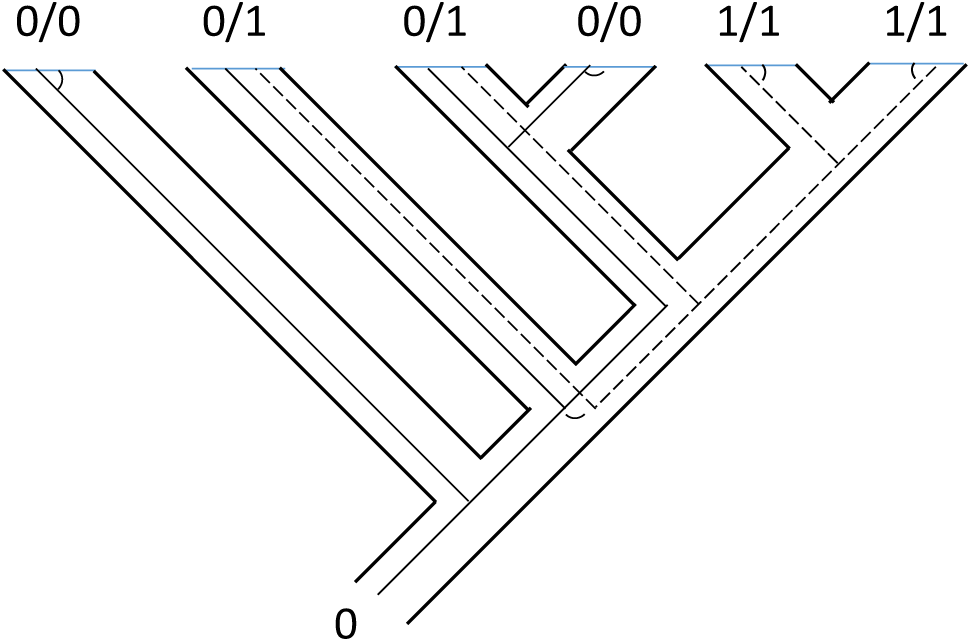
Genotype tree, Ψ, containing the optimal imbedded allele tree, *t*_*min*_. Tree *t*_*min*_ is constructed from the two minimal subtrees for the two alleles, 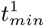 (dashed line) and 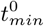 (fine solid line). These two trees join at a coalescence (duplication) event indicated by a small curved edge. Note the coalescence of homozygotes is shown by very short curved edges near the leaves of Ψ. A deep coalescence on Ψ is an edge containing “extra edges” of the imbedded allele tree for its full length. There are four such edges on Ψ, each having one extra imbedded edge. This tree has the fewest possible deep coalescences, among all allele trees with these genotypes that obey the infinite sites model. Its DC score, 4, is the same as its PP score (see Fig. 1), which is also equal to the number of “overlapping” edges of Ψ—those containing both allele subtrees. Note also that the edges showing “polymorphism” or “deep coalescence” under the two approaches are the same. See Appendix.

Given *t*_*min*_, the number of deep coalescences can be easily computed based on the subtree of Ψ in which the two subtrees of *t*_*min*_ overlap, *ψ* (Fig. 2):

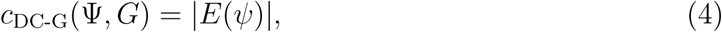

where |*E*(*ψ*)| is the number of edges of*ψ*.

The form of the optimal allele tree is of less interest than the fact that it leads to the following result:

#### Theorem 1.

*Given a rooted genotype tree*, Ψ, *and genotype character, G, and assuming that the pair of alleles underlying the genotypes are encoded by a binary character evolving on a rooted allele tree according to the infinite sites model (i.e., at most one* 0 → 1 *transition permitted, with no reversals), then*

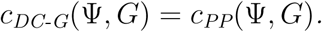

In other words, the PP score is the same as the DC score of an optimal imbedded allele tree for those genotypes. See Appendix for Proof. This also means that the DC-G problem can be solved in *O*(*n*) time, because it is equivalent to solving the PP problem.

### MDC-G: Inferring a genotype tree from genotype characters

The “Minimize Deep Coalescences” (MDC) problem, which infers a species tree from a collection of gene trees by minimizing the sum of DC scores across the gene trees (Maddison 1997; Ma et al. 2001; Maddison and Knowles 2006; Than and Nakhleh 2009; Bansal et al. 2010; Zhang 2011), can be modified to use genotypes alone.

### Problem (MDC-G Tree)

Given a collection of genotype characters, {*G*_*i*_}, find the genotype tree, 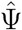, that minimizes

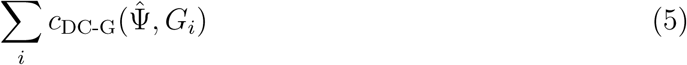

By Theorem 1, this is the same tree inferred by optimizing the PP criterion across these genotypes. Since an implementation of PP is available in Phylip (Felsenstein 2005), it is not necessary to implement the more involved computation implied by the definition of the DC-G score. The decision problem version of finding a tree using the PP score is NP-complete (Day and Sankoff 1987), and thus MDC-G is at least as computationally hard. However, heuristic search implementations benefit from the *O*(*n*) runtime of the inner loop score calculation in the PP algorithm (Felsenstein 1979).

## Materials and Methods II. Data Analysis

### Genome Sequence Data

We surveyed population genomic variation in the saguaro cactus (*Carnegiea gigantea* (Engelman.) Britton & Rose) using short read genome sequence data for 20 diploid individuals in 10 populations across its range. We used FASTQC v.0.11.8 (Andrews 2018) to examine all read sets and Trimmomatic 0.38 (Bolger et al. 2014) to trim low quality ends. Options for the latter were either “ILLUMINACLIP:TruSeq3-PE:2:30:10 HEADCROP:3 LEADING:20 TRAILING:20 SLIDINGWINDOW:4:20 MINLEN:36” or “ILLUMINACLIP:TruSeq3-PE:2:30:10 CROP:147 HEADCROP:3 LEADING:20 TRAILING:20 SLIDINGWINDOW:4:20 MINLEN:36” (for the Orégano and San Marcial samples), depending on quality control reports. Trimmed reads were mapped to the saguaro SGP5 v.1.3 assembly (“SGP5” henceforth) (Copetti et al. 2017) using Bowtie2 v. 2.3.4.3 (Langmead and Salzberg 2012) in paired end mode, discarding unaligned, mixed and discordant pairs. Deduplication was implemented with Samtools v. 1.9. (Li et al. 2009) with the command sequence (sort, fixmate, sort, markup -r) described in their documentation, and resulting BAM files were indexed with samtools. Statistics for resulting BAM files were obtained using the samtools stats command, and coverage was computed based on a SGP5 genome size of 1.403 Gbp from the C-value database (Bennett and Leitch 2012).

Subsets of sites in three genomic regions were extracted from BAM files based on the SGP5 assembly. The first consisted of all 1893 scaffolds larger than 100 kb, which totalled 281 MB, or about 1/4 of the assembled genome (“Large100k” data set). The second and third subsets comprised zero-fold and four-fold degenerate sites in codons of all protein coding genes (“0-Fold” and “4-Fold” data sets), which were extracted using a custom PERL script (adapted from “identify_4D_sites.pl”, see https://github.com/tsackton/linked-selection/blob/master/misc_scripts/Identify_4D_Sites.pl). The SGP5 annotation v.1.3 was used as input.

Genotypes were called from the 20 BAM files with bcftools (options: mpileup -q 20 -Q 20 -a ‘FORMAT/DP’), followed by bcftools (Li et al. 2009) (options: call -m) to create files with all reads. The FORMAT option allows downstream filtering by individual and site depth. Each of these files were then hard filtered by bcftools (options: -filter –SnpGap 3 -i ‘%QUAL>20 && TYPE != “indel”‘). Then VCF files were normalized with bcftools (options: norm -d all) and checked again for duplicates with a custom PERL script. Further filtering by coverage depth and missing genotype calls was effected with a custom PERL script, VCFParse.pl. In general, a genotype call was required to have coverage of between 2-100x to be considered non-missing. Analyses in the pooled sample of 20 were required to have 13 of 20 genotype calls present at a site. The individual population samples were each required to have two of two genotypes present to keep a site. These were obtained by using bcftools (option: view -S) to subsample the VCF file for the entire data set.

### Genetic diversity

Genetic diversity estimators were obtained using two protocols. First, we used the genotype calls derived from the bcftools pipeline described above and calculated Tajima’s (*π*_*T*_) and Watterson’s nucleotide diversity estimator (*π*_*W*_) with our VCFParse.pl script. Watterson’s estimator was corrected for missing genotypes using the formula in Ferretti et al. (2012). Missing individuals at a site were omitted in the *π*_*T*_ calculation.

Second, we used the Angsd package (Korneliussen et al. 2014) on only the Large100k data to estimate these parameters directly from the BAM files. This approach sidesteps genotype calling and works directly at the level of genotype likelihoods. Angsd was run with settings chosen to match those used the bcftools pipeline as closely as possible, including hard filters to require “minQ 20 minMapQ 1 minIndDepth 2” and “minInd” set appropriately. Because ancestral states of SNPs were unavailable, folded site frequency spectra were used. BAM files for the species-wide 10 population sample consisted of the original 20 BAM files restricted to the appropriate genomic regions, but the 2-individual analyses for each population separately were done using just the two BAM files for that population. This differed from the genotype calling strategy using bcftools, in which the smaller samples were obtained by subsetting the genotype calls in the large sample.

The spatial distribution of genomic diversity was interpolated using point kriging with no drifts with a linear variogram and no nugget effect as recommended for small, noisy samples. The grid was generated using Surfer (Golden Software, Golden, Colorado, USA) overlaid on a shaded relief base map derived from GMTED2010 (https://topotools.cr.usgs.gov/gmted_viewer/viewer.htm).

### Phylogenetic Analysis

#### Complete data sets

“Complete” phylogenetic data matrices for all 20 individuals were prepared for each of the three genomic subsamples, and then these were thinned to account for statistical nonindependence between sites (see below). All sites in which genotype calls from the bcftools pipeline displayed exactly two alleles among the individuals in the sample were included in phylogenetic data sets. Data sets were formatted for use in Phylip (Felsenstein 2005) and PAUP* (Swofford 2002).

#### Phylogenetic non-independence across the genome

The equivalence of the PP and MDC-G score summed across multiple sites holds when sites are independent, so that each site can theoretically evolve on its own allele tree. If a set of nearby sites are linked as haplotypes evolving on the same allele tree, then the two scores could differ. Complete data sets should therefore be “thinned” to sites that are approximately independent of one another (Lee et al. 2014). We estimated the rate of decay of statistical dependence between sites using two procedures. In the first, we used PLINK v. 1.9 (Chang et al. 2015) to extract sites in approximate linkage equilibrium using a stringent requirement of 1.0 for the VIF parameter (option: –indep 50 5 1).

In the second procedure, we implemented a direct phylogenetic assay based on character compatibility (Felsenstein 2004) and a generalization of Hudson and Kaplan’s (1985) “four-gamete test” for a pair of unphased diploid genotypes. A pair of sites for *n* individuals is *pairwise genotype compatible* if and only if there exist 2*n* haplotypes evolving on some “perfect phylogeny” (one with no homoplasy). This is what is expected under the infinite sites model. Wang (2013) used this concept to identify blocks of compatible sites, but did not explicitly describe it; for completeness, we show the computation in the Appendix. For a sample of 10000 regularly spaced sites in each of the three data sets, we computed the fraction of sites at lag distance of *λ* sites downstream (1 ⩽ *λ* ⩽ 100) that are pairwise genotype compatible. This provides an estimate of autocorrelation as a function of coordinate distance. We then assigned a *thinning distance, k*_thin_, by inspection from these plots, defined as the minimum distance at which the pairwise compatibility fraction decreases to its genome-wide level.

#### Phylogenetic data set and tree construction

For a complete data set, *D*, of length *m* sites, we built *k*_thin_ thinned data sets, *D*_*i*_, of sites spaced evenly at *k*_thin_ intervals:

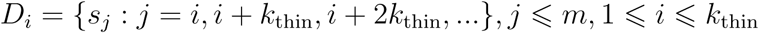

For each *D*_*i*_, we estimated a rooted genotype tree, 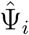, using the PP tree search implemented in dollop in Phylip 3.695 (2013) with settings of ‘Polymorphism parsimony’, ‘Ancestral’ states set to ‘?’, and 100 replicated random addition sequences using Phylip’s ‘Jumble’ option. Prior to search, the taxon order in each matrix was also randomized by a PERL script. A bug in the program’s output of the total PP score was fixed (Felsenstein, pers. comm.). Because dollop is not optimized for searching for the best *rooting* per se (Felsenstein, pers. comm.), the optimal tree reported by dollop was rerooted in all possible ways using a PERL wrapper script around the Newick Utilities program nw reroot (Junier and Zdobnov 2010), and each rerooted tree was scored with dollop as described above. The best rooted tree was retained.

For a single thinned data set, such as those generated by running PLINK on each of our complete data sets, we estimated standard bootstrap support using Phylip’s seqboot in conjunction with dollop as described. However, for the data sets constructed from the compatibility-based thinning, which produced *multiple* data sets subsampled from each complete data set, we constructed an overall estimate of the tree by computing a (rooted) majority rule consensus of these:

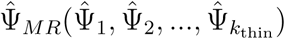

However, trees in 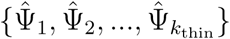 may not be independent of each other, because the sites used to construct, say, 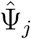 are not independent of the sites used to construct 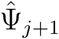. In fact, the sites at position *i* in these two data sets are only one character apart in the original complete matrix. We computed a measure of autocorrelation of the estimated trees as a function of distance between sites in *D* (“lag”) using pairwise tree-to-tree rooted Robinson-Foulds distances. This was only sensible for the Large100k data set, which has a sizable maximum lag distance of *k*_thin_ = 100. For each lag distance *λ* : 1 ⩽ *λ* ⩽ *k*_thin_/2, we computed 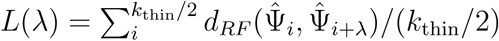, where *d*_*RF*_ (Ψ_*i*_, Ψ_*j*_) is the rooted Robinson-Foulds distance between trees Ψ_*i*_ and Ψ_*j*_.

Because PP trees consistently estimated the root to be near population samples that had higher missing data than average, we performed a numerical experiment to check for the influence of low coverage individuals on the method. For each of the three complete data sets, we downsampled sites by preferentially keeping sites that were present in the three individuals with the most missing data, until the overall percentage of missing data in those three samples was decreased below 5-8%, which was equivalent to levels found in several other samples. Resulting data sets were about 1/3 the size of the originals. We then reran the thinning and PP searches as described above, but changed the *k*_thin_ values to maintain independence but correct for the smaller sizes of these matrices.

Finally, we compared our results with trees obtained with SVDquartets (Chifman and Kubatko 2014) in PAUP*, a scalable phylogenomic inference method. To force SVDquartets to treat individuals as genotypes, each individual was recoded as two sequences, randomly phasing all heterozygous positions, and then each pair was forced to be treated as a “species” under the multi-species coalesent assumption of the algorithm. Optimal trees from replicate thinned data sets were combined as described above for PP analyses, with random phasing done separately for each thinned data set.

## Results

### Sequence Data and Genetic Diversity

We obtained a total of 198 Gbp of short read sequence for 20 individuals in 10 populations (Table 1). Average coverage ranged from 2.2x to 17.5x (mean = 7.6x). The San Marcial coverage was the lowest by population.

**Table 1.** Sequencing read statistics for samples

| Sample | Population | Reads ($10^6$) | Bp ($10^9$) | Coverage (x) | Mean Length (bp) | Mean Quality Score |
| --- | --- | --- | --- | --- | --- | --- |
| Sample_C60 | Caborca | 87.18 | 11.64 | 8.3 | 133 | 33 |
| Sample_C61 | Caborca | 92.95 | 12.28 | 8.8 | 132 | 33 |
| Sample_D17 | El Dipo | 65.93 | 6.26 | 4.5 | 94 | 37 |
| Sample_D73 | El Dipo | 74.01 | 7.05 | 5.0 | 95 | 37 |
| Sample_G50 | Guasimas | 83.68 | 11.19 | 8.0 | 133 | 33 |
| Sample_G81 | Guasimas | 81.44 | 10.90 | 7.8 | 133 | 33 |
| Sample_K26 | Kino Bay | 87.85 | 11.78 | 8.4 | 134 | 33 |
| Sample_K4 | Kino Bay | 56.04 | 7.44 | 5.3 | 132 | 33 |
| Sample_J34 | La Joyita | 59.52 | 5.67 | 4.0 | 95 | 37 |
| Sample_J45 | La Joyita | 65.96 | 6.29 | 4.5 | 95 | 37 |
| Sample_M69 | Masiaca | 71.93 | 6.84 | 4.9 | 95 | 37 |
| Sample_M85 | Masiaca | 43.39 | 4.12 | 2.9 | 94 | 37 |
| SGP-OR3 | Oregano | 89.82 | 11.69 | 8.3 | 130 | 34 |
| SGP-OR6 | Oregano | 76.94 | 9.99 | 7.1 | 129 | 34 |
| SGP-SM1 | San Marcial | 28.90 | 3.78 | 2.7 | 130 | 34 |
| SGP-SM7 | San Marcial | 22.98 | 3.03 | 2.2 | 131 | 34 |
| SGP5-L004 | Tucson | 256.20 | 24.49 | 17.5 | 95 | 38 |
| Sample_CU | Tucson | 76.08 | 7.24 | 5.2 | 95 | 37 |
| Sample_119 | Wickenburg | 110.12 | 10.48 | 7.5 | 95 | 37 |
| Sample_129 | Wickenburg | 73.01 | 6.96 | 5.0 | 95 | 37 |

The bcftools pipeline inferred from just under 50,000 to nearly 3 million variants in the three data sets (Table 2). Almost all were biallelic: the fraction of variants having 3 or 4 alleles ranged from only 0.0035 - 0.0045.

**Table 2.** Variants, genotype statistics and phylogenetic data sets

| Data set | 0-Fold Sites | 4-Fold Sites | Large 100k Scaffolds |
| --- | --- | --- | --- |
| Total effective sites | 19563072 | 4592714 | 267653586 |
| Number of variants | 106998 | 47741 | 2871509 |
| Biallelic sites in complete matrix | 106629 | 47511 | 2858685 |
| Tri- or quad-allelic sites | 369 | 230 | 12824 |
| Fraction of tri- or quad-allelic sites | 0.0035 | 0.0048 | 0.0045 |
| Fraction of those sites homozygous | 0.72 | 0.72 | 0.74 |
| Fraction of those sites heterozygous | 0.19 | 0.19 | 0.17 |
| Fraction of those sites missing | 0.08 | 0.09 | 0.09 |
| Fraction of those sites with no missing data | 0.18 | 0.17 | 0.14 |
| Sites in each compatibility-thinned matrix | 5332 | 4752 | 28586 |
| Sites in PLINK-thinned matrix | 27457 | 13977 | 88480 |
| Sites in downsampled matrix | 31994 | 14367 | 755194 |
| Sites in each thinned downsampled matrix | 5333 | 4789 | 7552 |

Table 3 summarizes all nucleotide diversity estimates obtained with the bcftools pipeline, subsets of the genome and different populations. An overall estimate of neutral genetic diversity based on 4-fold degenerate sites in protein coding genes is 0.0025, which is quite low compared to many plant species. The ratio of 0-fold to 4-fold diversity is quite high, around 50%, indicating a high proportion of nonsynonymous diversity. Tajima’s and Watterson’s estimators of nucleotide diversity are in broad agreement, as are the estimates obtained from the bcftools pipeline vs. the Angsd pipeline, which uses genotype likelihoods directly (Table 4).

**Table 3.**
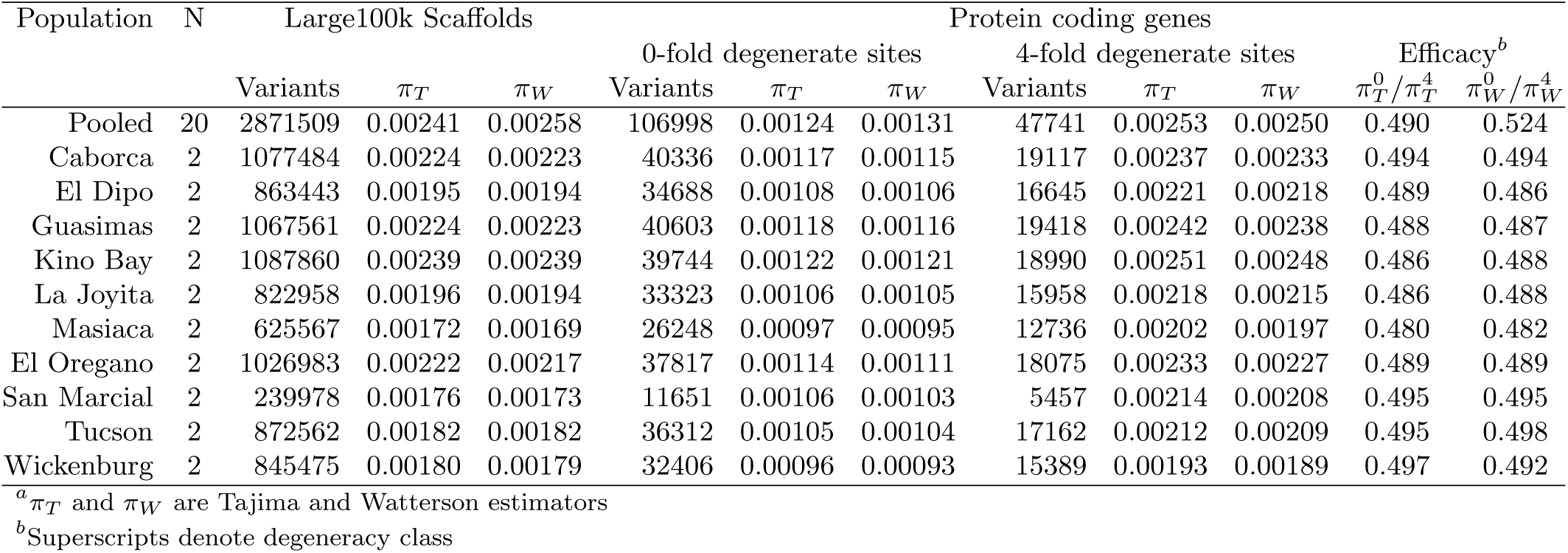
Nucleotide diversity estimates^a^

**Table 4.** Nucleotide diversity estimates (Angsd)

| Population | N | Large100k | Scaffolds |
| --- | --- | --- | --- |
| | | $\pi_T$ | $\pi_W$ |
| 10 pops(x2) | 20 | 0.00277 | 0.00325 |
| Caborca | 2 | 0.00235 | 0.00232 |
| El Dipo | 2 | 0.00203 | 0.00199 |
| Guasimas | 2 | 0.00243 | 0.00240 |
| Kino Bay | 2 | 0.00255 | 0.00253 |
| La Joyita | 2 | 0.00198 | 0.00194 |
| Masiaca | 2 | 0.00181 | 0.00174 |
| El Oregano | 2 | 0.00250 | 0.00245 |
| San Marcial | 2 | 0.00180 | 0.00173 |
| Tucson | 2 | 0.00184 | 0.00183 |
| Wickenburg | 2 | 0.00185 | 0.00181 |

A geographic perspective on genetic diversity can be seen in Figure 3. The highest diversity is around Kino Bay in Sonora (Table 3), but it drops off relatively quickly to the east and south, and more slowly to the north. Masiaca and Wickenburg, with generally the lowest diversities, are populations near the southern and northern distribution margins respectively. The spatial distribution is similar for both estimators of nucleotide diversity. Using the Watterson estimator for the 4-fold degenerate sites in protein coding, the isoline of 0.0021 delimits well what we consider saguaro populations in good health and number. Although the interpolated surface is only derived from the genomic data, there is a significant negative linear trend with elevation (Fig. 3), and with the vegetation units described by Shreve (1951), displaying the highest genomic diversity in the Central Gulf Coast, intermediate values in the Arizona Upland and the thornscrub, and the lowest diversity in the transition to the Mojave Desert.

**Fig. 3.**
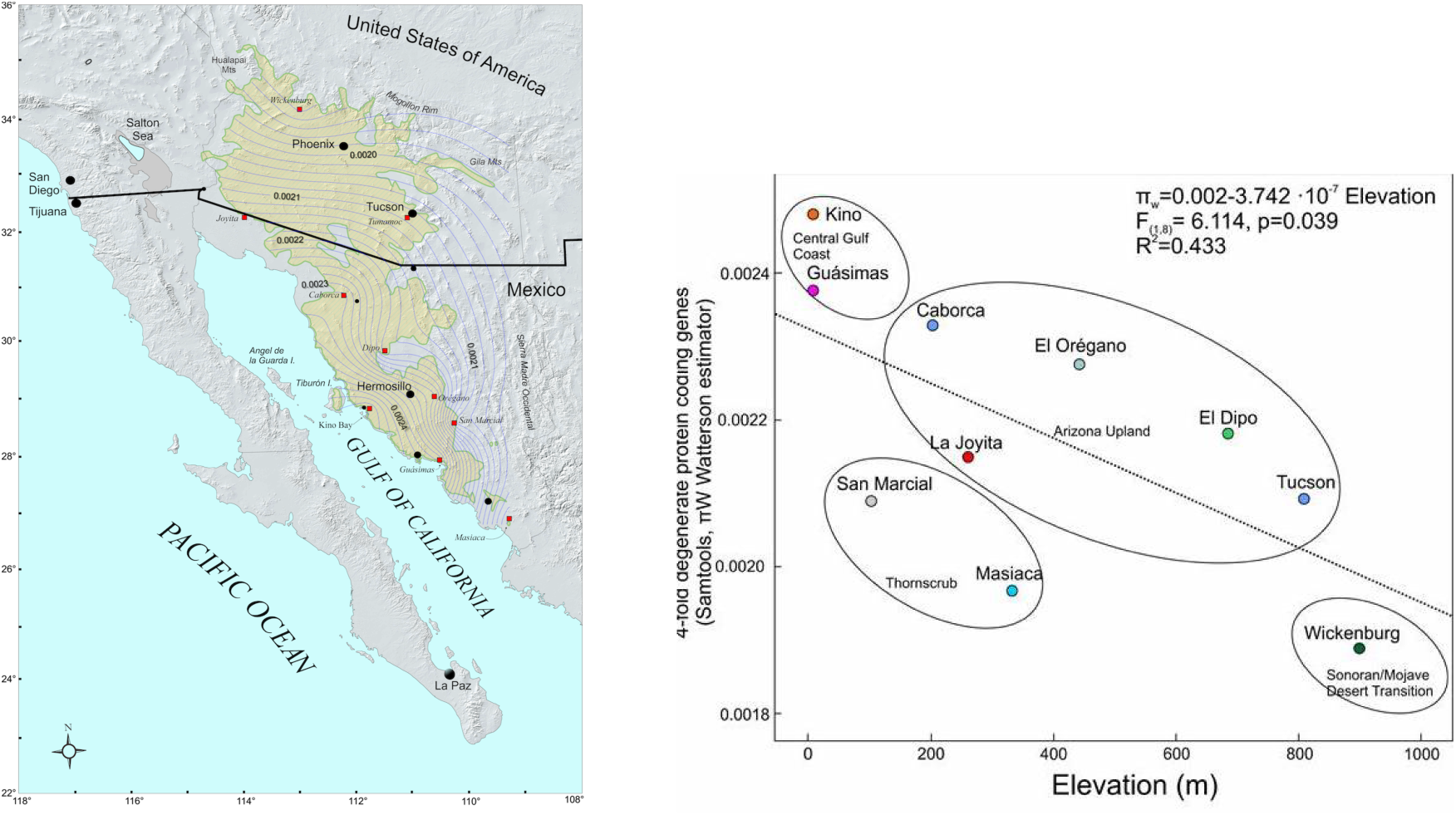
a) Contour plot of estimated nucleotide diversity (*π*_*W*_)in 4-fold degenerate sites in 10 populations of saguaro. Range of saguaro shown in yellow. b) Relationship between elevation and estimated nucleotide diversity (*π*_*W*_).

### Phylogenetic data sets

The three complete phylogenetic data sets ranged in size from 47511 and 106629 sites for 4-fold and 0-fold degenerate sites to nearly 3 million in the Large100k data set (Table 2). The fraction of genotypes called as heterozygous ranged from 17-20%, whereas the fraction of sites with missing data was 8-9%. Missing data was distributed unevenly among samples, with the two San Marcial samples having 40-54% missing genotypes and the Masiaca M85 individual having 20%.

Analyses of pairwise genotype compatibility in the three complete data sets revealed a strong signal of local statistical dependence along the genome coordinate axis, presumably due to linkage (Fig. 4). The 4-fold data set lost dependence most quickly, within a *k*_thin_ of 10 sites; the 0-fold data was about 20 sites; and the Large100k data about 100 sites. Note that these distances are in units of sites in the data matrices: two neighboring sites in the phylogenetic data set might be separated by a long coordinate distance along the scaffold because of intervening nonvariable sites.

**Fig. 4.**
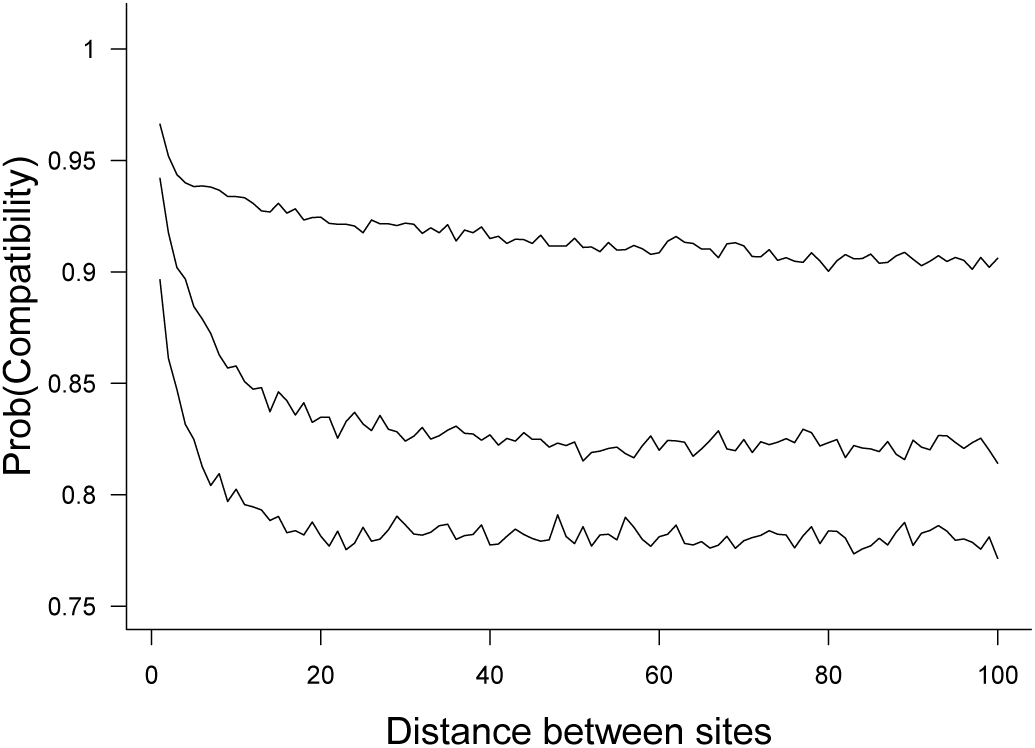
Decay of pairwise genotype compatibility versus distance between sites in data sets (from top to bottom: Large100k; 0-fold; 4-fold).

Thinning by PLINK’s algorithm for identifying sites in approximate linkage equilibrium produced three data sets having 27457, 13977, and 88480 sites for 0-fold, 4-fold, and Large100k data sets, respectively (Table 2). These all correspond to average thin distances that are smaller than found by our pairwise compatibility method and thus produced data sets with more sites each.

### Intraspecific Genotype Phylogenies

Trees inferred using PP and compatibility thinning in the three data sets were highly congruent (Fig. 5). A northern clade of four populations, Caborca, La Joyita, Tucson and Wickenburg, was consistently strongly supported, though La Joyita and Caborca within this clade showed slightly lowered support for the monophyly of individual populations. The remaining populations were all highly supported as monophyletic except at the root. Rerooting of the dollop solution with our rerooting script often improved the optimality score, but the root was consistently still placed within the San Marcial population. The Kino Bay population, which has the highest nucleotide diversity and lies at sea level on the Sea of Cortez to the west of San Marcial was nested further within the genotype phylogeny. Trees inferred from the Large100k data differed slightly with respect to Kino Bay, having it as sib group to the rest of the tree, except for San Marcial.

**Fig. 5.**
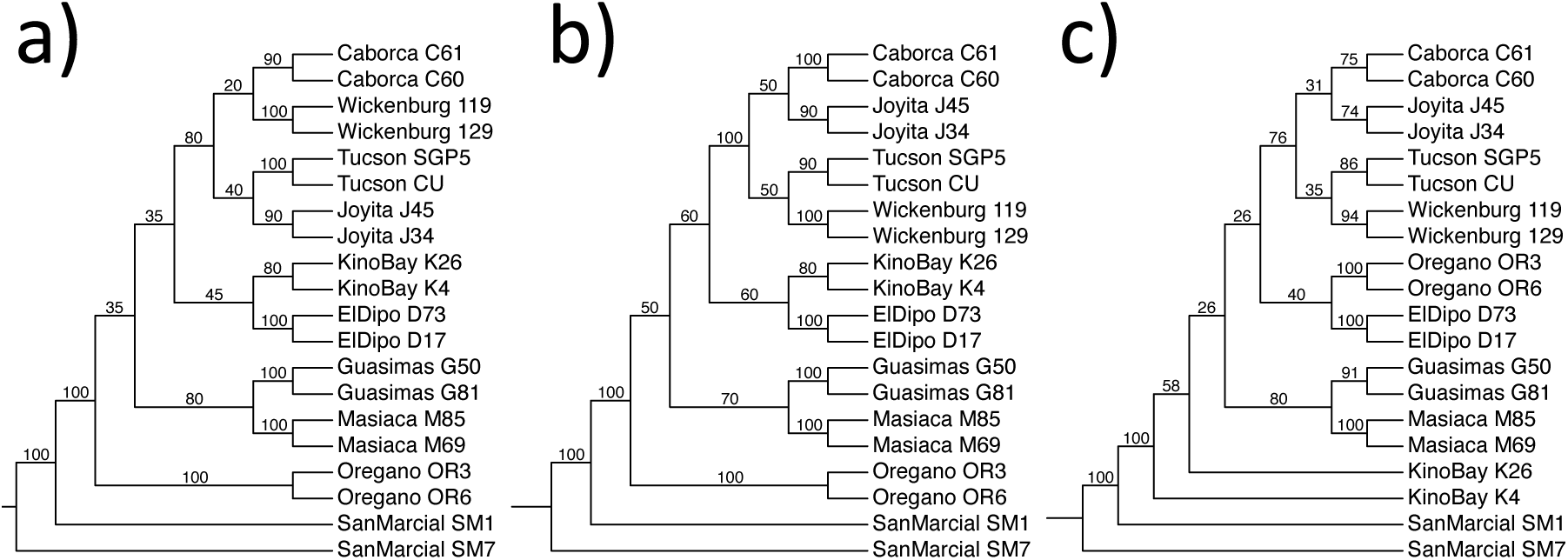
Genotype trees reconstructed with PP in Phylip. These are majority rule trees of sets of *k*_thin_ compatibility-thinned data sets for a) biallelic 0-fold sites (*k*_thin_ = 20); b) biallelic 4-fold sites (*k*_thin_ = 10); c) Large100k sites (*k*_thin_ = 100); Proportions on edges reflect the number of trees containing indicated bipartitions. Trees are rooted by the PP method.

Figure 6 assesses the autocorrelation between trees inferred by PP from the compatibility-thinned data sets for the Large100k data. The Robinson-Foulds distance between trees estimated at different lag distances appears to show no relationship relative genome coordinate distance, implying that there is no autocorrelation among the PP trees inferred from the thinned data sets.

**Fig. 6.**
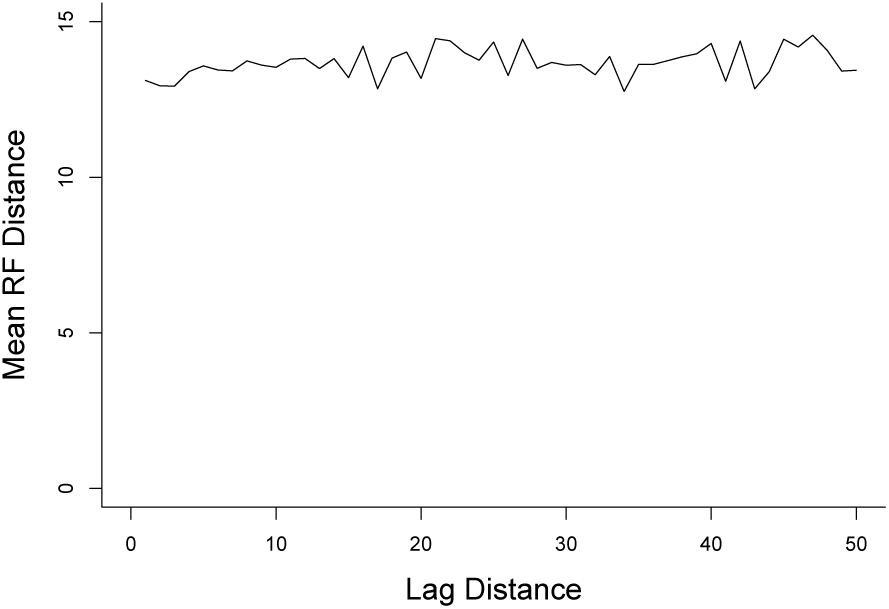
Mean RF distances between trees estimated from *k*_thin_ = 100 thinned data sets subsampled from Large100k data set, 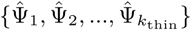, at different lag distances. Lag distance, *λ*, refers to a pair of trees, 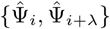.

Trees inferred with PP using the data sets thinned via PLINK (Fig. 7) all agreed with the 0-fold and 4-fold trees from PP and compatibility thinning, and also were rooted in San Marcial.

**Fig. 7.**
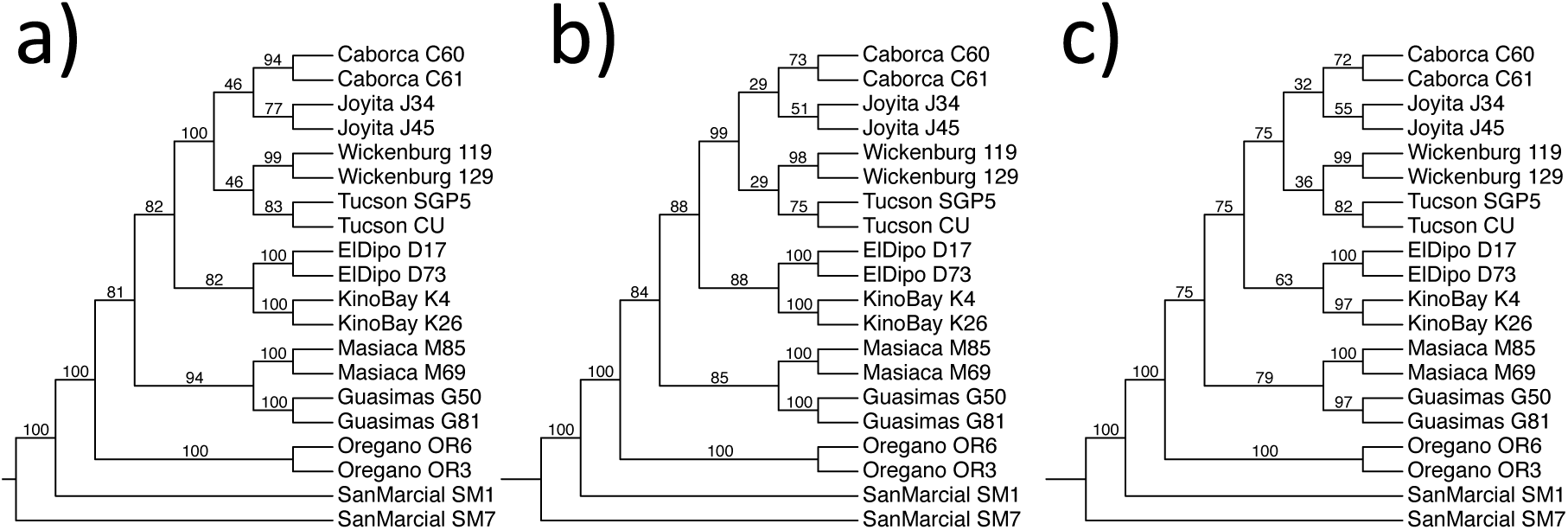
Genotype trees reconstructed with PP in Phylip. These are bootstrap majority rule trees from the three data sets thinned to approximate linkage equilibrium in PLINK for a) biallelic 0-fold sites; b) biallelic 4-fold sites; c) Large100k sites. Trees are rooted by the PP method.

Downsampling the three data sets to assess the impact of missing sites resulted in smaller data sets (Table 2), but with much more even distributions of missing data (Table 5). Because of their smaller size, the 0-fold and 4-fold downsamples were thinned to fewer data sets: 6 and 3 data sets respectively, which maintains approximately the same thinned data set size as in the non-downsampled data. The Large100k data was thinned into 100 data sets (a *k*_thin_ value that should produce sites thinned well beyond the *k*_thin_ distance in the original Large100k data set). The 0-fold and 4-fold majority rule trees (Fig. 8) were identical to those in Fig. 5. However, the Large100k analysis departed from this otherwise consistent picture, albeit with only moderate bootstrap values. It was rooted in the Masiaca population at the southern limit of the saguaro range.

**Table 5.** Fractional missing data in downsampled datasets

| Sample | 0-Fold Sites | 4-Fold Sites | Large100k |
| --- | --- | --- | --- |
| Caborca.C60 | 0.0092 | 0.0094 | 0.0054 |
| Caborca.C61 | 0.0160 | 0.0144 | 0.0078 |
| ElDipo.D17 | 0.0553 | 0.0575 | 0.0544 |
| ElDipo.D73 | 0.0298 | 0.0289 | 0.0395 |
| Guasimas.G50 | 0.0148 | 0.0123 | 0.0094 |
| Guasimas.G81 | 0.0161 | 0.0144 | 0.0109 |
| Joyita.J34 | 0.0728 | 0.0770 | 0.0850 |
| Joyita.J45 | 0.0366 | 0.0380 | 0.0536 |
| KinoBay.K26 | 0.0144 | 0.0107 | 0.0081 |
| KinoBay.K4 | 0.0640 | 0.0611 | 0.0500 |
| Masiaca.M69 | 0.0400 | 0.0397 | 0.0506 |
| Masiaca.M85 | 0.0519 | 0.0717 | 0.0588 |
| Oregano.OR3 | 0.0155 | 0.0174 | 0.0115 |
| Oregano.OR6 | 0.0262 | 0.0264 | 0.0196 |
| SanMarcial.SM1 | 0.0297 | 0.0301 | 0.0594 |
| SanMarcial.SM7 | 0.0416 | 0.0409 | 0.0542 |
| Tucson.CU | 0.0284 | 0.0298 | 0.0314 |
| Tucson.SGP5 | 0.0051 | 0.0036 | 0.0004 |
| Wickenburg_119 | 0.0090 | 0.0061 | 0.0074 |
| Wickenburg_129 | 0.0308 | 0.0322 | 0.0337 |

**Fig. 8.**
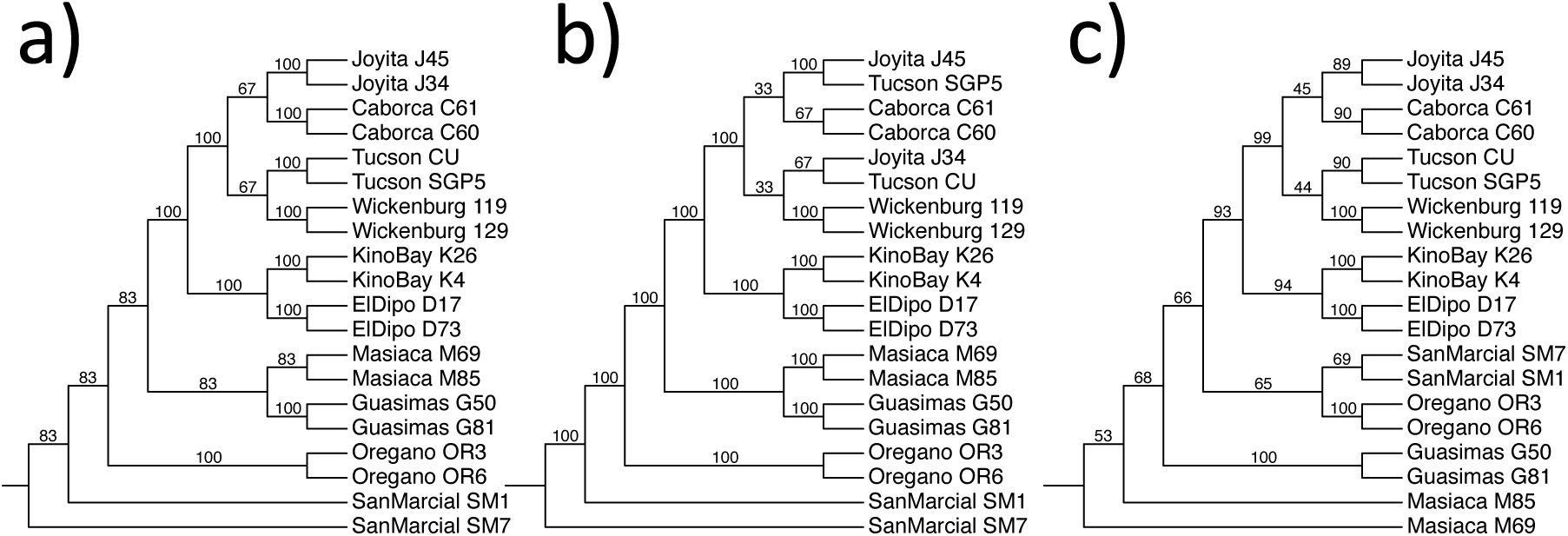
PP genotype trees based on downsampled data sets (see Table 5). Trees are majority rule trees of sets of *k*_thin_ thinned data sets for a) biallelic 0-fold sites (*k*_thin_ = 6); b) biallelic 4-fold sites (*k*_thin_ = 3); c) Large100k sites (*k*_thin_ = 100); Proportions on edges reflect the number of trees containing indicated bipartitions. Trees are rooted by the PP method.

Comparable trees constructed by SVDquartets are shown in Figure 9 for the two gene data sets. The unrooted topology is highly congruent between 0-fold and 4-fold data sets; all pairs of individuals within populations are supported as clades; and there remains some variability in the relationship of the northernmost populations. One minor difference with the PP trees is that SVDquartets moves the San Marcial population by one nearest neighbor interchange so it is slightly further from Orégano on the tree..

**Fig. 9.**
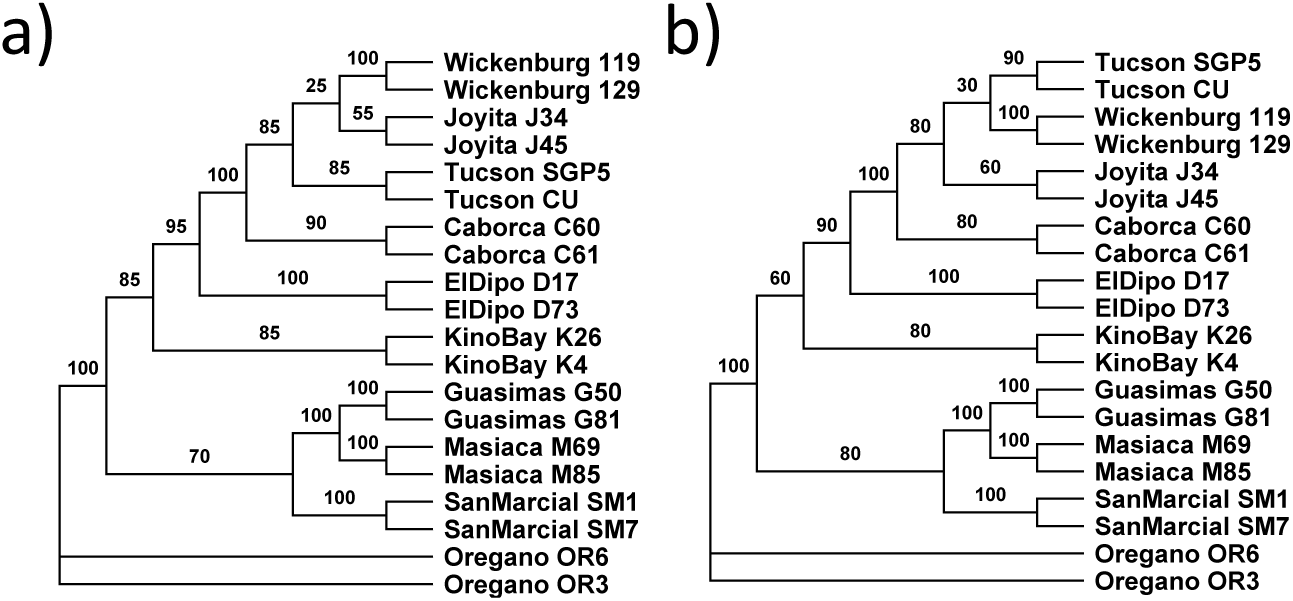
Genotype phylogenies based on SVDquartets thinned by compatibility thinning as in PP analyses. Phylogenies are based on a) 0-fold, and b) 4-fold data sets. Proportions on edges reflect the number of trees among the thinned runs containing indicated bipartitions. Trees are unrooted.

## Discussion

Inger (1967) published a phylogeny of frogs based on musculo-skeletal characters using code written by Felsenstein, implementing polymorphism parsimony for the first time (Felsenstein 1979). Felsenstein (1979) and Farris (1978) later formalized the problem statements and algorithms for assigning a PP score to a tree, both citing for inspiration work on *Drosophila* inversion polymorphisms. The algorithm presented by Felsenstein is similar in spirit and runtime to other parsimony scoring methods and shares with them the potential to scale to large data sets, at least with heuristic tree search algorithms (though, like MP and ML, PP is NP-complete: Day and Sankoff 1987). Despite the prevalence of polymorphic character data in the real world, and the obvious connection to diploid genotype data, PP has been used sparingly: e.g., for plant morphological data (Baum 1983), and bird retrotransposons (Suh et al. 2015).

In this paper we showed that polymorphism parsimony can be placed in the framework of minimizing deep coalescences in the genotypes’ underlying allele tree. This is intuitive on the one hand—the language of minimizing the extent of polymorphism on a tree is similar to minimizing deep coalescences—but also somewhat surprising. One method takes only discrete character data as input, while the other seems much more directly related to a reconciliation between an allele tree and genotype tree. The criterion of minimizing the number of polymorphic edges in the genotype tree is equivalent to minimizing deep coalescences in the “best” allele tree that can be constructed from the genotype data. An equivalent interpretation of these optimality criteria is that they minimize the number of “fixation/loss” events (Farris 1978), that is, losses of one or the other allele from a polymorphic ancestor.

Though intuitive, this equivalence is not automatic. It holds if the allele tree evolves according to an infinite sites model. In this setting the equivalence of the two methods places PP from the late 1970s into a rich framework of gene tree reconciliation, which dates to about the same time (Goodman et al. 1979) but has led to a very large computational, algorithmic and empirical literature (Page 1994; Maddison 1997; Nakhleh 2013).

One particularly attractive feature of polymorphism parsimony is that it provides an estimate of the root, because its underlying asymmetrical character evolution model penalizes rootings with deeper polymorphisms. For example, the tree,

(0/0, (0/0, (0/1, 0/1))) has a PP score of two, but the rerooted tree, (0/1, (0/1, (0/0, 0/0))) has a score of three. In the saguaro data this provided some evidence to dispute the hypothesis that the population with highest nucleotide diversity in saguaro is also at the root of the tree. PP shares this feature with some long standing methods, such as midpoint rooting, UPGMA and likelihood inference using a molecular clock (Huelsenbeck et al. 2002; Felsenstein 2004), and some more recent methods of species tree inference, such as BPP (Rannala and Yang 2017).

### Assumptions

The PP method applied to genome-wide genotype data is likely to be most appropriate when individual sampling is sparse relative to population differentiation and isolation. For example, one individual sampled per population within a species in which population differentiation is significant may balance the requirement of limited gene flow, so that an individual is a stand-in for a population, with limited sequence divergence, so that the infinite sites assumption is met. As sampling within panmictic populations increases, we expect more unresolved nodes within populations on the genotype tree, and lower confidence levels.

In the saguaro data we sampled two individuals per species, which allowed tests of monophyly of the 10 populations. Had their been little phylogenetic structure between populations, these support values should have been low, but most populations were supported at 90% levels or above for a variety of data sets and treatments. The main exception was the rooting of the trees within the San Marcial population, which causes this population to be paraphyletic in several analyses.

The algorithm for minimizing deep coalescence forces homozygous alleles in an individual to coalesce at the leaves of the genotype tree. This is what a simple minimization criterion based on the DC score should do, and it is engineered by the reconcilation mapping function (Yu et al. 2011, Thm 6), but it is unlikely to be accurate for individuals that are closely related within a single population. Under the coalescent model every labeled history on the 2*n* alleles is equally probable (Xu and Yang 2016), and the chance of alleles of a genotype character pairing up as siblings within each individual on the allele tree is small. The parsimony rationale is to assume shallow coalescence unless there is evidence of deep coalescence based on the genotype data, but a prior based on the Kingman coalescent and random mating would disagree. This naturally places a premium on answering the basic question of how much the sampled individuals depart from random mating.

In general, the validity of an infinite sites model for our data is difficult to test, because homoplasy in the data can be caused by either multiple hits or discordant gene trees (hemiplasy: Avise and Robinson 2008) or both. The fraction of tri- or quadri-allelic variants was less than 0.5% in our three data sets, which implies that the rate of multiple hits is low.

### Thinning

Thinning SNP data to a subset of sites that are in linkage equilibrium and phylogenetically independent is recommended for a number of methods of SNP-based phylogenetic inference, including PP, SVDquartets and variants (Chifman and Kubatko 2014; Vachaspati and Warnow 2018), and SNAPP (Bryant et al. 2012). We used a method implemented in PLINK based on the correlation between genotype vectors at pairs of sites, and a new method based on pairwise phylogenetic compatibility, to thin data. Even though we used a high stringency in the PLINK analysis, it tended to retain more sites than the phylogenetic assay. The resulting genotype trees on these larger data sets were highly congruent with our results from compatibility-thinned data, and both sets of results agreed about the root of the saguaro genotype phylogeny.

These results raise several questions for future work. First, as SNP data sets for phylogenetic inference grow more dense, the size of thinned data sets will only increase to a certain point, and the question of how to combine these data sets will have to be addressed. Here we showed that there was little correlation between the trees constructed from the different thinned data sets, and we proposed a simple consensus tree to serve as a summary statistic, but it remains somewhat surprising that these trees are uncorrelated when there is strong evidence that the underlying site data are correlated locally in the genome. Second, once data sets are thinned enough, there remains the question of how many thinned data sets to construct: fewer with more sites, or more with fewer sites. Finally, there remains an open problem of how to identify true coalescent independent sites rigorously in the face of recombination, sequencing error, and homoplasy.

### Haplotype based inference

An alternative to thinning is to infer haplotype blocks from all the SNPs (reviewed in Browning and Browning 2011; Gusfield 2014). In a local block of nonrecombining sites, all SNPs should evolve on the same haplotype tree, which motivates the Perfect Phylogeny Haplotype Problem (Gusfield 2002). For short tracts and under the infinite sites assumption, all SNPs should be compatible with one haplotype tree, or nearly so. In practice, sequencing error and homoplasy can limit the applicability of this approach. More generally, haplotype trees that are not perfect phylogenies can be sought (Sridhar et al. 2007), but the search space gets quite large. Not only is there the usual search space across trees, but there is the additional exponential growth in the number of alternative haplotypes for a given set of heterozygous SNPs. An additional empirical problem evident in our saguaro data is that relatively low genetic variation can mean that there is a long coordinate distance between SNPs, which leads to a tradeoff between combining enough SNPs to build a reliable haplotype tree before reaching recombination distances that allow independent coalescent genealogies (Springer and Gatesy 2016). A general problem is to find a set of intervals that either partition or cover the SNP sites within which there is a perfect haplotype tree, but between which the trees may differ (Gramm et al. 2009; Wang 2013). Given a collection of such haplotype trees across genomes for multiple individuals, we would still have the problem of how to integrate them into a genotype tree. The classical MDC problem, using individuals as leaves of the containing tree and a set of independent haplotype trees, would be one potential solution.

### Relationship to other methods

Because the inner loop of tree search heuristics for PP involves a linear time PP scoring algorithm (Felsenstein 1979), runtimes for our data were quite managable even for the largest data sets examined here, some of which (after thinning in various ways) had nearly 100,000 sites. No runs took more than about one hour of CPU time on an HPC linux node, and the bootstrap and thinning replicates could be trivially distributed on the cluster to make the analyses described here quite tractable.

Other computationally fast methods for inferring a population tree that scale to data sets of the size considered here include TreeMix (Pickrell and Pritchard 2012), which constructs a covariance matrix based on gene frequencies from SNPs and infers a tree on which migration events can then be overlain; and SVDquartets (Chifman and Kubatko 2014; Vachaspati and Warnow 2018), which uses a non-parametric statistic based on the multi-species coalescent for quartets and then combines the quartets to build a full tree. The former assumes a simple allele frequency diffusion model and the latter a standard DNA substitution model of the kind used in molecular phylogenetics, together with the MSC. Both methods infer an unrooted tree. TreeMix can summarize genotype data as allele frequencies within populations, but SVDquartets needs sequences to be entered for each allele and then populations to be circumscribed. RevPoMo (Schrempf et al. 2016, 2019) is analogous to an allele frequency approach but models transitions between allele count states instead. It returns an unrooted tree though a predecessor (De Maio et al. 2015) returned a rooted tree. Its authors “discourage … using revPoMo on sequence data where no population data is available yet” (Schrempf et al. 2016, p. 369) because of the need for demographic parameters in the model.

Perhaps the most computationally ambitious current methods, BPP (Rannala and Yang 2017), IMA-3 (Hey et al. 2018) and *Beast2 (Ogilvie et al. 2017), use a model-based Bayesian approach to integrate over population genetics, demography and phylogenetics. However, these appear to come at the cost of perhaps two or more orders of magnitude slower running times (Hey et al. 2018; Ogilvie et al. 2017). These methods can return a rooted tree by assuming a clock or the infinite sites assumption itself. Polymorphism parsimony provides a rapid alternative for estimating a reasonable genotype or population tree first, which can then be used in a two-step procedure to make more nuanced inferences about demography, admixture and migration.

### Saguaro nucleotide diversity and phylogeography

Estimates of nucleotide diversity for saguaro as a whole were near 0.0025, which is low among plants even compared to some other long lived trees such as spruce, which has about 2.5 times the diversity of saguaro (Chen et al. 2019). The efficacy of selection (Chen et al. 2017), defined as the ratio of 0-fold to 4-fold degenerate site diversities, was nearly 0.49, which is also unusually high among plants (c.f., 0.44 in one spruce species, Chen et al. 2019). This indicates much higher relative genetic diversity for potentially fitness-related variants than seen in other plants. These results may stem from low effective population sizes and/or historical population crashes during glacial/interglacial periods. Irrespective of cause, the 0-fold phylogenetic data set we analyzed here had twice as many sites as the 4-fold degenerate data set and evidently as much or more information about genotype phylogeny.

Genetic diversity was highest at sea level at Kino Bay, Sonora, Mexico, a bit south of the approximate center of the species’ range. Diversity dropped off quickly to the south and east and more slowly to the north. The range of saguaro has been heavily influenced by the ebb and flow of North American glaciation. Evidence from packrat midden remains indicates, for example, that saguaros were absent in the US until recent reinvasions about 10,000 years ago (McAuliffe and Van Devender 1998). Given that the species split from its closest relatives several million years ago (Copetti et al. 2017), it seems likely that glacial refugia must have existed somewhere in the south of its current range or perhaps further south.

Our PP analyses places the root of the saguaro genotype tree to the east or possibly south of Kino Bay in either the San Marcial or Masiaca populations, which are at the margins of the current range of saguaro. The latter two populations were favored differentially by analyses that were thinned in different ways, suggesting that such thinning choices can be significant. None of the analyses directly supported a rooting within the Kino Bay population. San Marcial and Masiaca are in thornscrub vegetation and both have substantially lower genetic diversity than Kino Bay.

Few simple genetic patterns are evident in the geographic distribution of saguaro and other columnar cacti (Bustamante et al. 2016). This is particularly true for columnar cacti in their northwestern range of distribution where frost limits these sensitive plants. In the case of the saguaro, its range is constrained by elevation (towards the Sierra Madre Occidental to the east and the Mogollon Rim to the north), westward by the Gulf of California, and most likely by the very low precipitation of the Gran Desierto where Sonora, California and Arizona meet (Albuquerque et al. 2018).

The pattern of higher diversity in the center of saguaro’s range around Kino Bay does generally support the so-called “center-periphery hypothesis” (Pironon et al. 2016) but likely this is modulated by dynamic changes in the very recent past (Lázaro-Nogal et al. 2017) and details of the demography and evolution of these populations. The position of the southern populations of San Marcial and Masiaca as outgroups to the remaining populations hints at possible crashes during the interglacial aggravated by ongoing global climate change. These factors likely led to differentiation and genomic diversity reduction through small effective population size and restricted local gene flow. Albuquerque et al. (2018) found that the saguaro distribution is contracting. They estimated a mean loss of almost 7% by 2050 under different climate change scenarios. That contraction is occurring on the western edge of the range from western Arizona to the southernmost extent in Mexico. Our rooted trees largely imply a basal grade of small, isolated southern and southeastern populations giving rise to several distinct large, thriving populations to the north and the northwest, which reinforces the idea of a refuge near the southern boundaries of the actual distribution range at the lowest elevations. The actual distribution of extant nucleotide diversity suggests a Pleistocene refuge in the lowlands of the Gulf of California coast in southern Sonora, and the small range of diversity estimates may indicate a rapid expansion northwards during the interglacial. If we exclude the outlier populations of San Marcial and Masiaca and concentrate on more central, numerically abundant, and/or the expanding, continuous populations in the northern range, there is a hint of a latitudinal decrease in nucleotide diversity. Further analyses on correlations between SNP variation and climatic, geographic and ecological variables are necessary to understand these issues more fully.

## Acknowledgements

We thank Joe Felsenstein for rapidly tracking down a bug in Phylip’s dollop program and providing a fix.

## Appendix

We assume each tree, *T*, is rooted, with edge set, *E*(*T*), node set, *V* (*T*), and leaf set, *L*(*T*) ⊂ *V* (*T*). Define the internal node set 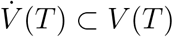 to be all nodes with (in or out)degree two or more. Each leaf *x* ∈ *L*(*T*) has indegree one and is labeled from the set *χ*. Let *𝒯*_*χ*_ be the set of all such rooted trees. An internal node *v* with outdegree one is called *unary* and two is *binary*. If all 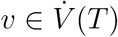 are binary, *T* is *binary*.

For *T* ∈ *𝒯*_*χ*_, the most ancestral node having outdegree two or more, if it exists, is called the *crown root, ρ*_*T*_. A node with indegree zero and outdegree one, if it exists, is called the *stem root*, 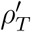. If *T* has a crown root with indegree zero it is called a *crown tree*. If *T* has a stem root that is the parent of a crown root node, then *T* is a *stem tree*, with a stem edge, 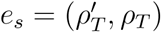 (see Fig. 10).

**Fig. 10.**
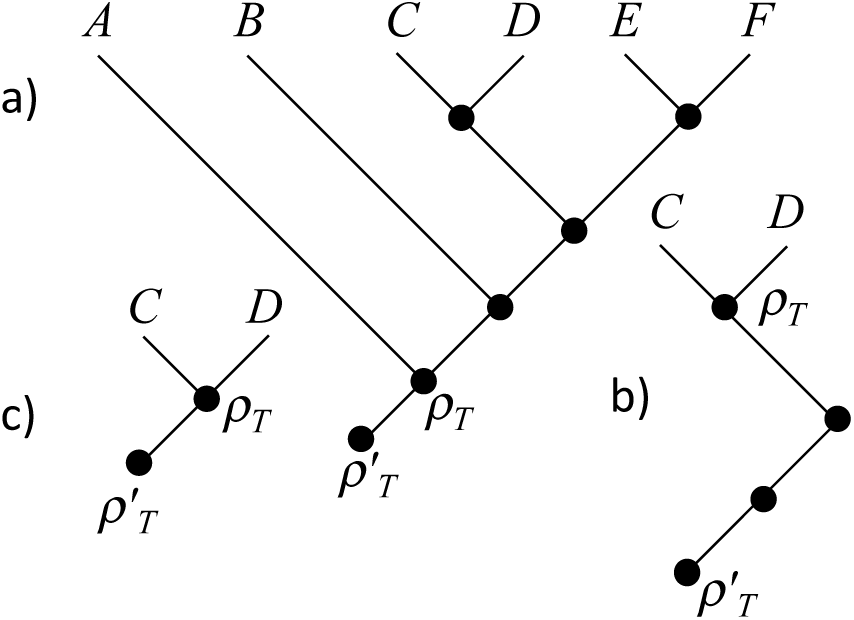
Illustration of rooted stem tree, *T*, and two subtree operations. Let 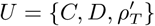 a) Original rooted stem tree, (note notation for stem and crown root nodes); b) subtree *T* ‖_*U*_ (note retention of unary nodes), which is neither a stem tree or crown tree by our definitions; c) subtree *T* |_*U*_ (note suppression of unary nodes), which is a stem tree.

If node *v*′ is an ancestor of *v*, we write *v*′ > *v*. Define *d*_*T*_ (*v*′, *v*) as the number of edges on the path between *v*′ and *v*, where *d*_*T*_ (*v, v*) = 0.

The set of leaves descended from *v* is *C*_*T*_ (*v*). The node of the most recent common ancestor of a set of leaves, *A*, is MRCA_*T*_ (*A*). For any binary 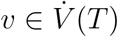, let *l*(*v*), *r*(*v*) and *a*(*v*) be the left child, right child and parent node of *v*, respectively, if these exist.

Define *T* ‖_*U*_ to be the minimal subtree of *T* containing a set of nodes *U* ⊆ *V* (*T*), and *T* |_*U*_ is obtained from *T* ‖_*U*_ by suppressing any unary nodes on *T* ‖_*U*_ (Ma et al. 2001) (See Fig. 10).

### Solution to PP Score Problem (Felsenstein 1979)

For reference, we describe Felsenstein’s (1979) two-pass algorithm to reconstruct ancestral states under polymorphism parsimony, modified slightly to allow missing data. Let the genotype tree, Ψ be a rooted binary stem tree, and denote the genotype states of a biallelic site, in which alleles are labeled 0 or 1, with an ordered pair of boolean variables to indicate their presence or absence (*y*^0^, *y*^1^), *y* ∈ {+, −}. Thus, genotypes 0/0, 0/1, 1/1, and ./. correspond to (+, −), (++), (−+), (−, −), respectively.

More generally, let 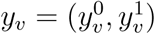 be a pair of boolean variables that indicates the presence or absence of the respective alleles among any descendant leaves of *v*. If *v* is a leaf node, this is exactly the genotype of *v*. The algorithm first does a downpass over 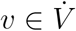, assigning *y*_*v*_ as follows:

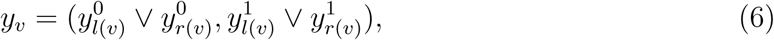

where present/absent is treated as true/false for the logical “or” operator, ∨.

Next an uppass traversal over 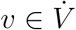 computes the actual optimal ancestral genotype states:

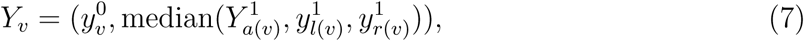

where “median(*b*_1_, *b*_2_, *b*_3_)”, is the most frequent state in three boolean variables. At *v* = *ρ*_Ψ_, the state 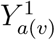 is initialized to ‘−’ because 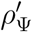 has genotype of (+, −), by assumption. The asymmetry in the two components of *Y*_*v*_ arises because, by assumption, the 0 allele is always present in the ancestor of the root, but the 1 allele is not. Once the final internal states have been inferred, then the PP score, *c*_*P P*_ (Ψ, *G*), is the number of nodes for which *Y*_*v*_ = *Y*_*a*(*v*)_ = (+, +).

### Solution to DC-G Score Problem

#### Preliminaries

In addition to the rooted binary stem genotype tree, Ψ, also now assume an underlying diploid rooted binary stem allele tree, *t*, for which there are at most two alleles for each leaf of Ψ. For any set of leaves of *t, W* ⊆ *L*(*t*), let *α*(*W*) be the corresponding set of leaves of Ψ.

Define the *MRCA-mapping, ℳ*, from node *v* in *t* to a node in Ψ as *ℳ*_Ψ_(*v*) = MRCA_Ψ_(*α*(*C*_*t*_(*v*))).

Define a *coalescent history* as a mapping, *h*, of all nodes in *t* to nodes in Ψ. Informally, this describes how the allele tree is imbedded in the genotype tree. For edge, *e* = (*u, v*) ∈ *E*(*t*), let *D*_Ψ_(*e, h*(*t*)) = *d*_Ψ_(*h*(*u*), *h*(*v*)) be the number of edges in Ψ in the coalescent history’s path from *h*(*u*) to *h*(*v*) (Fig. 11). Now, the *optimal coalescent history, h*^*\**^(*t*), minimizes

**Fig. 11.**
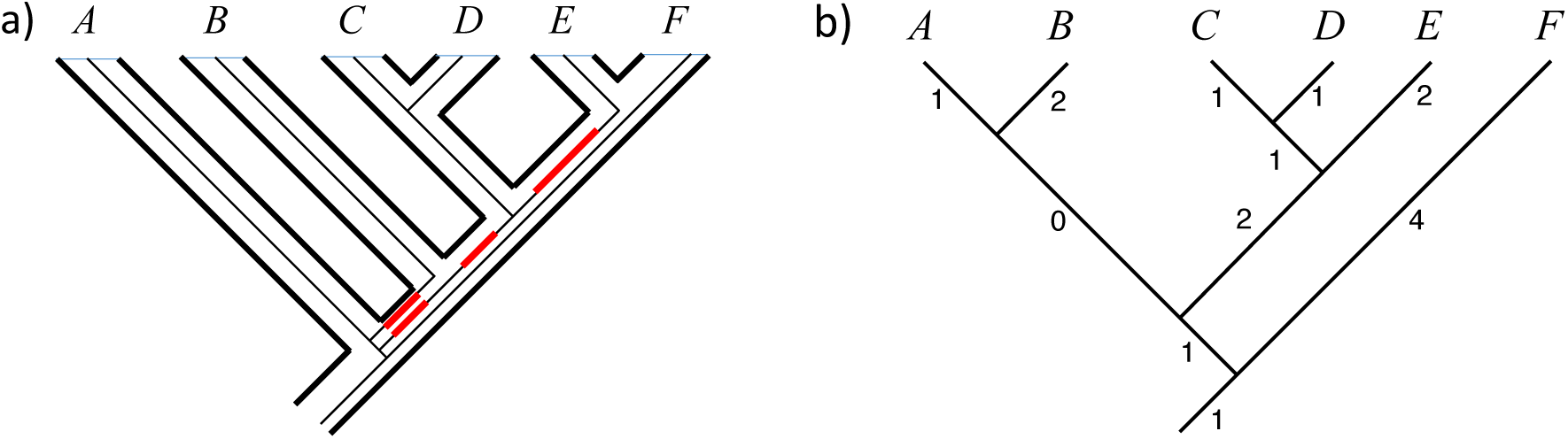
a) Genotype tree, Ψ, with imbedded allele tree, *t*; extra lineages shown in red; b) Allele tree, *t*. Numbers next to edges are *D*Ψ(*e, h*(*t*)), which count the number of edges in Ψ in which this edge of *t* is imbedded. Here |Ψ‖_*L*(*t*)_ = 11 from (a), ∑ _*e*_ *D*_Ψ_(*e, h*(*t*)) = 15 from (b), and DC score is therefore 4.

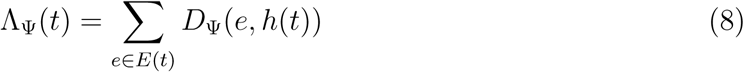

over all *h*(*t*). Although the number of distinct histories, *h*(*t*), can be quite large (Degnan and Salter 2005), the optimal history for a crown tree is just given by *h*^*\**^(*v*) = *ℳ*_Ψ_(*v*) for all 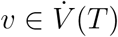 (Yu et al. 2011; Bayzid and Warnow 2012). For a stem allele tree, however, we need the following.

##### Lemma 1

(Optimal coalescent history for stem allele tree). *Assume g is a stem allele tree with crown tree, g*_*c*_, *and stem edge* 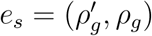, *and let u in* Ψ *be the node to which ρ*_*g*_ *maps via ℳ. Further, constrain* 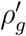 *to be present at some node, u*′ ⩾ *u of* Ψ *so that there is a path on* Ψ *between u*′ *and u having d*(*u’ u*) ⩾ 0 *edges. The optimal coalescent history of g has two parts: that for g*_*c*_, *which is determined by ℳ, as with any crown allele tree, and the history below this node down to u*′, *comprising a single edge of g. Thus*, Λ_Ψ_ (*g*) = Λ_Ψ_(*g*_*c*_) + *d*(*u*′, *u*).

*Proof.* The optimal coalescent history of *g*_*c*_ is given by *ℳ* (Yu et al. 2011), which maps *ρ*_*g*_ to *u* on Ψ. With *e*_*s*_ present, 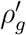 is forced to be at *u*′ ⩾ *u*, corresponding to *d*(*u*′, *u*) edges of Ψ between *u*′ and *u*. By the definition of *ℳ*, the only way to modify the coalescent history of *g* would be to move *ρ*_*g*_ closer to the root of Ψ, decreasing *d*(*u*′, *u*). However, because *ρ*_*g*_ has outdegree two, this would pull two allele tree lineages deeper in the genotype tree, and increase Λ_Ψ_(*g*_*c*_) by *two* for each edge lost in *d*(*u*′, *u*). Thus, the optimal coalescent history of *g* is given by the history for *g*_*c*_ plus the path corresponding to *e*_*s*_. □

There is some ambiguity in the literature (Than and Nakhleh 2010; Yu et al. 2011; Bayzid and Warnow 2012, 2018) (and in software implementations: Bayzid and Warnow 2012) about counting imbedded allele tree lineages and computing the DC score when the allele tree is missing from some leaves of the genotype tree: i.e., when *α*(*L*(*t*)) ⊂ *L*(Ψ). In that case, *t* could be considered to be imbedded in either Ψ|_*L*(*t*)_ or in Ψ‖_*L*(*t*)_ (Zhang 2011; Bayzid and Warnow 2012), and since the number of edges differs, so too could the DC score. We adopt the framework established in Bayzid and Warnow (2012) and Bayzid and Warnow (2018) which uses Ψ‖_*L*(*t*)_. This counts edges of Ψ joined at unary nodes of Ψ‖_*L*(*t*)_, whereas Ψ|_*L*(*t*)_ collapses these into a single edge.

#### DC score for stem tree with missing data

Now we are in a position to define the deep coalescence score in more general terms. The DC score, *c*_DC_(Ψ, *t*), is the number of “extra” imbedded allele tree lineages above what is minimally necessary given the genotype data. Suppose leaf set *A* ⊂ *L*(Ψ) has genotype data present, and suppose the stem root of allele tree *t* is forced to be present at some node *u* on Ψ, then let *U* = *A* ∪ {*u*}. The minimum number of edges of Ψ that must have at least one imbedded allele tree edge is |*E*(Ψ‖_*U*_)|, and therefore

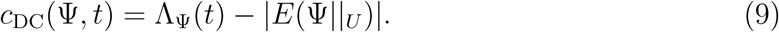

The last term depends only on the genotype tree and the data, not the unknown allele tree, *t*, so minimizing Λ_Ψ_(*t*) is equivalent to minimizing the number of deep coalescences.

#### Optimal allele trees

Now, we need to *find* an allele tree, *t* that minimizes Λ_Ψ_(*t*) for given Ψ over all *t* consistent with *G* and the infinite alleles assumption. If there were no constraints on *t*, the lemma below could be used to find this tree easily. However, *t* cannot be just any tree; it is constrained by the assumptions of the model so that: (i) it must include two disjoint subtrees, *t*^0^, and *t*^1^, the leaves of which have these respective allele states; (ii) the stem root node of *t*^0^ must be present at the stem root of Ψ, because the 0 allele is ancestral; and (iii) the stem edge of *t*^1^ (which is where the single 0 → 1 mutation occurs on *t*), attaches to some edge of *t*^0^. Denote the set of all allele trees satisfying conditions (i)-(iii) as 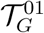.

Because Λ_Ψ_(*t*) is additive over edges, its value for a tree comprising two subtrees meeting at a single internal node is

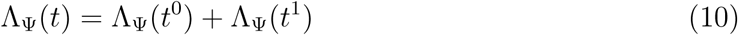

The following lemma shows how to find an optimal subtree for each of these terms separately. Because each of these terms has a lower bound determined only by the genotype data and Ψ, these optimal subtrees will minimize the sum, and therefore the assembled tree will also be the optimal.

##### Lemma 2

(Optimal allele tree). *Given a set of allele tree leaf nodes, W, with α*(*W*) *the corresponding leaves of* Ψ, *and u* = *ℳ*(*α*(*W*)), *and given a node, u*′ > *u of* Ψ, *at which the stem root of some allele tree, g, will be constrained to be present, then let U* = *α*(*W*) ∪ *u*′. *For all g* ∈ *𝒯*_*W*_, Λ_Ψ_(*g*) ⩾|*E*(Ψ‖_*U*_)|. *Moreover, there exists an optimal allele stem tree, g*_*min*_, *that achieves this lower bound, having the following properties: (i) for any leaf, x, of* Ψ, *having two leaves in W, the two leaves form a cherry (a pair of siblings) in g*_*min*_, *with a parent we call ξ*(*x*); *and (ii) the “backbone” subtree of g*_*min*_, *obtained by replacing any such cherries with their corresponding nodes, ξ*(*x*), *has a topology given by* Ψ|_*U*_.

*Proof.* Suppose *g*_*min*_ is given as above. Each node *ξ*(*x*) has two child edges, *e* = (*u, v*) in which both *u* and *v* map to node *x* on Ψ, because they correspond to the same genotype leaf, and thus *D*_Ψ_(*e, h*^*\**^(*g*_*min*_)) = 0 for both. Therefore, these cherries do not contribute to Λ_Ψ_(*g*).

Because the remaining “backbone” subtree, 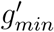, is the same as Ψ|_*U*_ (i.e., cluster by cluster), each edge of 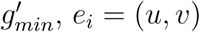, corresponds to a unique path, *p*_*i*_ in Ψ‖_*U*_ from *ℳ* (*u*) to *ℳ* (*v*). These paths together must partition Ψ‖_*U*_ edgewise and therefore Λ_Ψ_(*g*_*min*_) = |*E*(Ψ‖_*U*_)|. Alternatively, note that for every node of Ψ‖_*U*_ with outdegree two, *D*_Ψ_(*e*_*i*_, *h*^*\**^(*g*_*min*_)) = 1 (the path has one edge). For any node with outdegree one, the path has more than one edge, but in either case, these will be counted correctly in the value |*E*(Ψ‖_*U*_)|. □

#### Construction of t_min_

Now we describe the construction of *t*_*min*_, the optimal 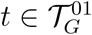. Let *W* ^0^ and *W* ^1^ be the subsets of leaf alleles having the 0 and 1 states respectively. The 0 allele must be present at the stem root of Ψ, by assumption, because it is ancestral according to the model. In Lemma 2 we therefore have 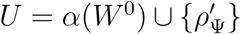, and 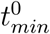 has the topology of Ψ|_*U*_ (see example in Fig. 10).

The solution for 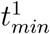 is complicated slightly by the final disposition of the stem edge of 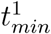, in particular, where it attaches to 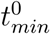. By this we mean also, because 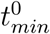 is imbedded at its optimal coalescent history, to which node, *u*″, in Ψ does it attach? The node 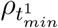 is placed at 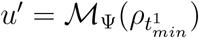, and according to Lemma 1, constraining 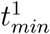 to be present at *u*″ adds *d*(*u*′, *u*″) to the score of imbedded allele edges. Thus we should choose the attachment point to minimize *d*(*u*′, *u*″). If 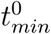 and *u*′ overlap, then we can choose *u*′ = *u*″. Then the stem root of 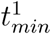 can attach to 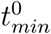 “just below” *u*′, effectively given the stem edge a length of zero in keeping with *d*(*u*′, *u*″) = 0. If they do not overlap, we choose *u*″ to be as close as possible to *u*′ in the path from *u*′ to the root, as long as *u*″ > *u*′.

This construction for *u*″ = *u*′ accords well with the reconcilation framework. For example, the MRCA mapping of both the stem and crown root of 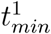 is *u*′, which means we infer a duplication at the stem root node, where it joins 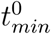 (Goodman et al. 1979). However, the “boundary” case of *u*″ > *u*′ is a bit more interesting, because it exposes some of the assumptions of the polymorphism parsimony model. This case arises only when there are no heterozygotes in the data and either all genotypes are trivially 0/0, or some genotypes are 1/1 and those genotypes form a clade on Ψ (if there is more than one 1/1). No internal node is reconstructed as 0/1 under these circumstances, but the PP model assumes that a 0/1 ancestor served as an intermediate. Moreover it also assumes a duplication/coalescent event was necessary to evolve the 0/1 genotype. Essentially, the model postulates a hidden polymorphic state in the tree despite none having been observed in the data.

We can engineer this hidden state within a reconciliation framework by inserting a node, *x* and pendant edge, *e*_*x*_, on the stem edge of 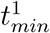. This subdivides the original stem edge into a parent edge, *e*_*a*_ and child edge, *e*_*c*_ that subtends the crown root node of 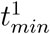 (Fig. 12). Node *x* represents the “duplication” event, and the origin of the new 1 allele state occurs along *e*_*c*_. The edge, *e*_*x*_ terminates immediately and represents the “loss” of the 0-allele. Because of this, *e*_*x*_ adds zero to Λ_Ψ_. Both allele trees are present simultaneously—briefly—indicating ephemeral polymorphism.

**Fig. 12.**
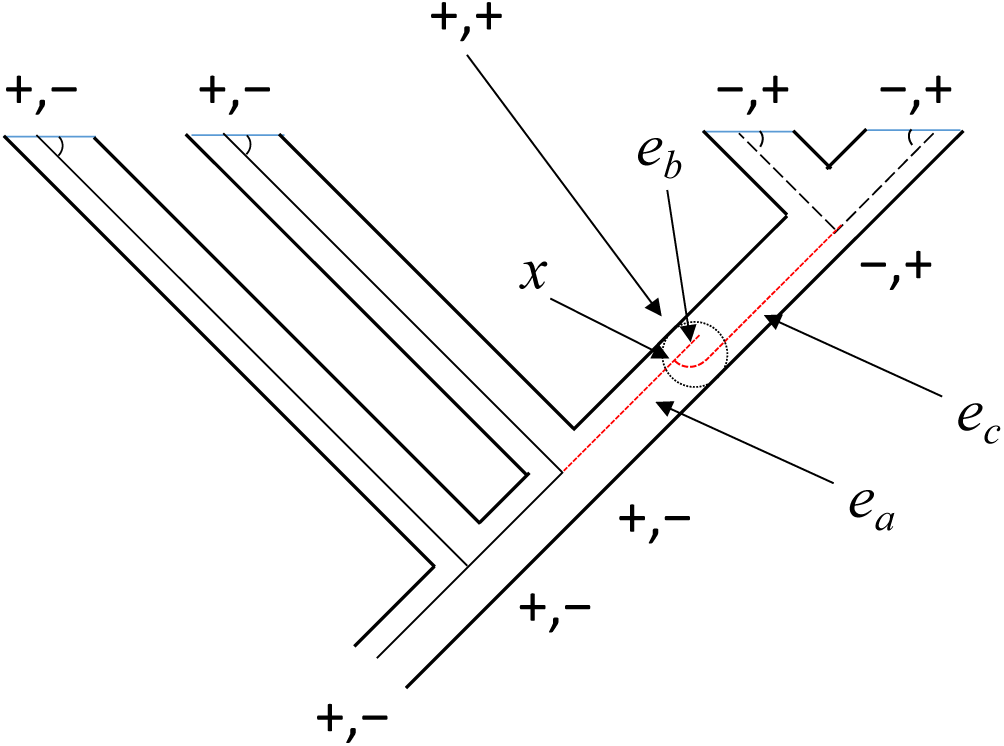
Illustration of a deeper interpretation for constructing *t*_*min*_, needed in the boundary case in which there is no overlap between allele subtrees, 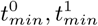. To maintain the assumptions of the PP model, we could insert a subtree (red dotted lines) inside the genotype tree edge in place of the stem edge of 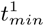. In this subtree there is a duplication, *x*, a mutation to 1 in one child edge, *e*_*c*_, and an immediate loss of the other edge, *e*_*b*_. This generates an ephemeral ancestral heterozygote (dotted circle).

In both cases, the construction of the two subtrees leads to an optimal allele tree with minimum DC score, but what is this score? Any edge of Ψ can have 0, 1, or 2 imbedded allele tree edges, but only edges with 2 contribute to the DC score. These are edges in which both 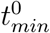 and 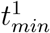 are present, which is the subtree of Ψ where the two subtrees overlap, *ψ*. Thus, the DC-G score is:

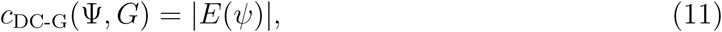

which makes clear that *c*_DC-G_(Ψ, *G*) is nonzero only if the two subtrees overlap.

### Proof of Theorem 1

We will show that the ancestral states reconstructed by 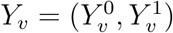, are equivalent to the presence or absence of alleles in an imbedded allele tree that matches *t*_*min*_ for the DC-G computation. Thus the two methods produce the same score for a given genotype tree and character.

The proof has two parts: (i) the PP downpass effectively traces out a tree for each allele that is the minimal subtree of Ψ for the leaves having that allele present and extending down to the crown root (for the 0 allele this will be the final tree, 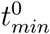; (ii) however, for the 1 allele, 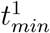 must be truncated toward the root so that it does not extend below the last possible point of origin of the 1 allele. This is taken care of by the uppass phase, which only affects the 1 allele.

To prove (i), note that for allele *i* ∈ {0, 1} at node *v*, the downpass value, 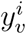 is set to + if any descendant nodes of *v* are +. The downpass thus traces out the minimal subtree of + states that extends from the crown root of Ψ to each leaf of Ψ having the allele. For the 0 allele this is the final set of states because the 0 allele is assumed to be present at the stem root of Ψ, so 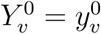, and the tree implied by PP is exactly 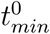 described above.

To prove (ii) is true for the 1 allele, consider the instances when the uppass might *change* a downpass value 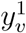. From Eq. 7, the uppass value, 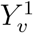, depends on states at its three adjacent nodes:

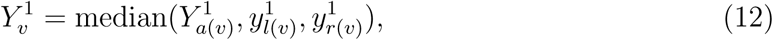

There are 8 possible combinations of + and − arguments in this function, and enumeration of these shows that only two of them lead to cases in which 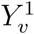 is different from 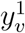 (partly because of how 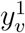 is computed from its two child states): either 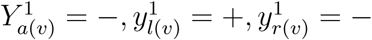 or 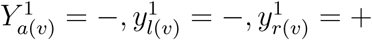. In both cases the change is from 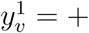 to 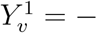. The uppass is initialized by setting the stem root state to − (see Eq. 7), so that 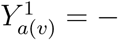 when *v* is the crown root. Thus, the uppass traversal *might* start out having to invoke one of these changes of state iff only one of its two children has the + state. If so, this will continue until the uppass reaches the most recent node that has both children with a + state; that is, a node that *must* have the 1 allele (rather than merely *may* have it).

This node of Ψ is then the proper location of the crown root node of 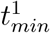, and the uppass is therefore equivalent to construction of the 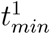 *s* crown allele tree. Its stem edge attaches to 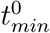 as described earlier. Since the DC-G score depends only on the two subtrees and their intersection (Eq. 11), this means that the PP score and DC-G score are the same (see Fig. 13). □

**Fig. 13.**
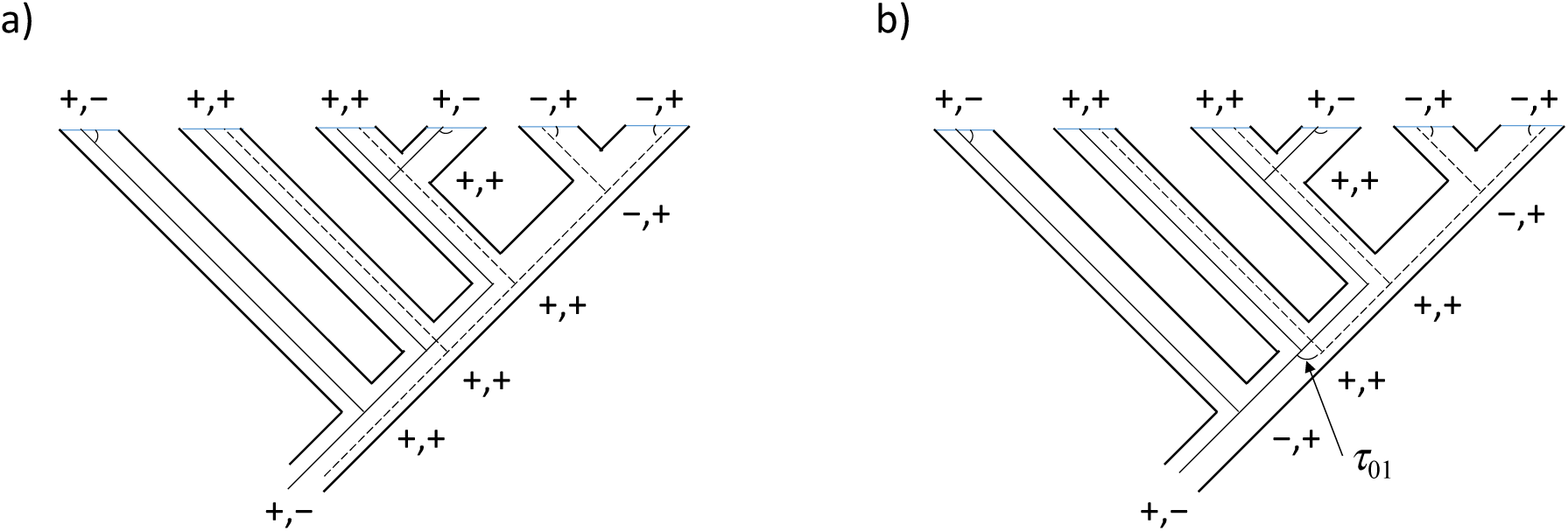
Example of equivalence between PP and DC-G ancestral state reconstructions. Solid line is allele tree 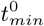; dashed line is 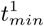. The stem edge of 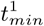 joins the two (here labeled *τ*01). a) After the downpass phase: initial allele subtrees for DC-G and the *y*_*v*_ states for each node are shown. b) After the uppass phase: final allele tree for DC-G and final PP ancestral states, *Y*_*v*_, are shown.

### Pairwise Genotype Compatibility

Under the infinite sites model in the absence of recombination there should be blocks of neighboring loci that are compatible with each other (Hudson and Kaplan 1985). Recombination and sequencing error limits the lengths of such blocks. To estimate the distance between effectively unlinked, statistically independent sites, we assayed compatibility among genotypes of sites. For example, suppose two sites’ genotypes for an individual are 0/1 and 1/1, then the possible haplotype assignments (= “gametes”) are 01 and 11, and each must be present. Thus, two of the four “gametes” (01 and 11) allowed by the four gamete test (Hudson and Kaplan 1985) must already be present in this one individual’s pair of genotypes. If the other two gamete types are also present in possible haplotype assignments for any other individuals in the data, then this pair of sites is genotype incompatible. The diagram in Fig. 14 is useful for enumerating the haplotype (=gamete) types that are implied by a pair of genotypes. By checking off observed gamete types among such genotype pairs for all individuals, one can determine if the two sites are genotype-compatible (see Wang 2013, for further details).

**Fig. 14.**
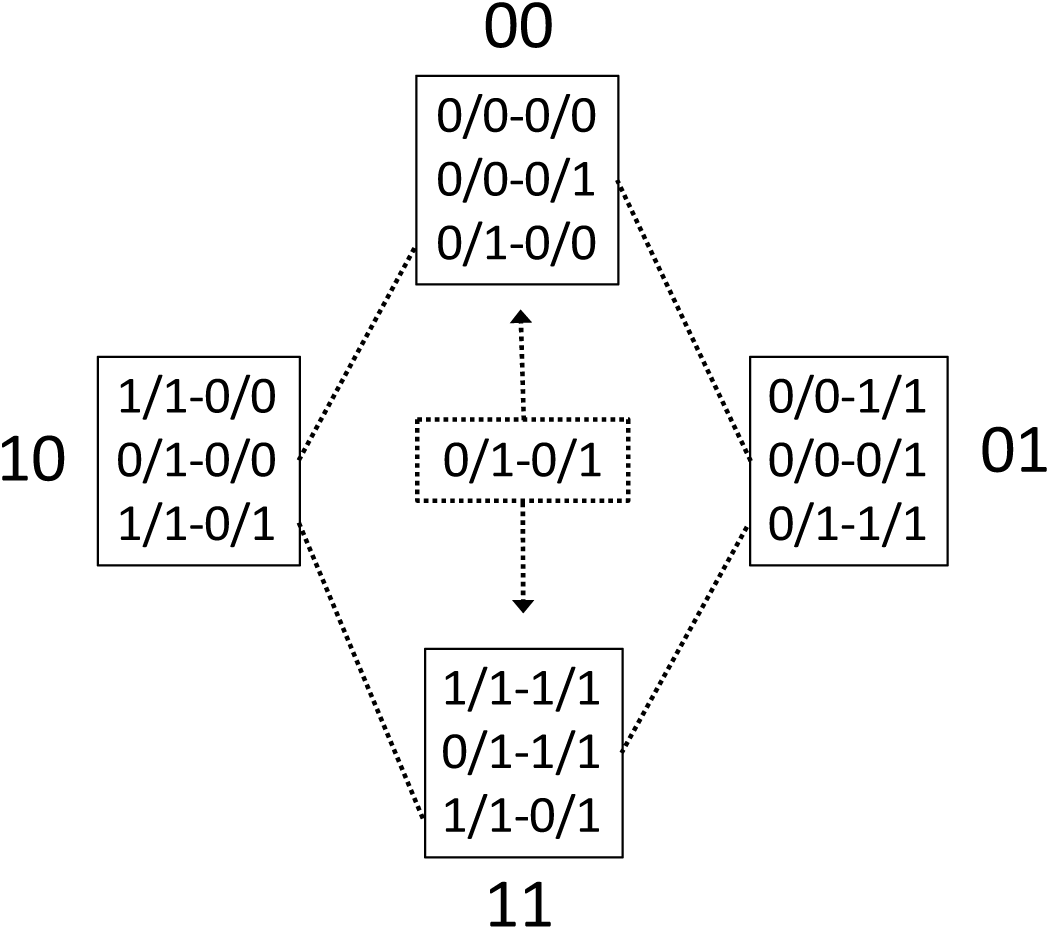
Pairwise genotype compatibility. Genotype pairs at two sites are shown within boxes. Large pairs of numbers are allelic haplotypes implied by the genotype pairs. In some cases (dashed connectors), a genotype pair implies both of two haplotype pairs. If individuals in the data set collectively have haplotypes (=“gametes”) of all four types, the pair of sites is “genotype-incompatible”. The 0/1-0/1 genotype pair is ambiguous and can go in either direction indicated, which means if three gamete types are already observed among the individuals, a 0/1-0/1 genotype pair will *not* cause genotype incompatibility.

This uses the unrooted definition of character compatibility so is an unrooted version of genotype compatibility.

## References

Albuquerque, F., B. Benito, M. Á.M. Rodriguez, and C. Gray. 2018. Potential changes in the distribution of *Carnegiea gigantea* under future scenarios. PeerJ 6:e5623.

Andrews, S. 2018. FastQC. Dowloaded from https://www.bioinformatics.babraham.ac.uk/projects/fastqc/.

Avise, J. C. and T. J. Robinson. 2008. Hemiplasy: A new term in the lexicon of phylogenetics. Systematic Biology 57:503–507.

Bansal, M. S., J. G. Burleigh, and O. Eulenstein. 2010. Efficient genome-scale phylogenetic analysis under the duplication-loss and deep coalescence cost models. BMC Bioinformatics 11:S42.

Baum, B. R. 1983. A phylogenetic analysis of the tribe Triticeae (Poaceae) based on morphological characters of the genera. Canadian Journal of Botany-Revue Canadienne De Botanique 61:518–535.

Bayzid, M. S. and T. Warnow. 2012. Estimating optimal species trees from incomplete gene trees under deep coalescence. Journal of Computational Biology 19:591–605.

Bayzid, M. S. and T. Warnow. 2018. Gene tree parsimony for incomplete gene trees: addressing true biological loss. Algorithms for Molecular Biology 13:1–12.

Bennett, M. and I. Leitch. 2012. Plant dna c-values database (release 6.0, dec. 2012).

Bolger, A. M., M. Lohse, and B. Usadel. 2014. Trimmomatic: a flexible trimmer for Illumina sequence data. Bioinformatics 30:2114–2120.

Bravo, G. A., A. Antonelli, C. D. Bacon, K. Bartoszek, M. P. K. Blom, S. Huynh, G. Jones, L. L. Knowles, S. Lamichhaney, T. Marcussen, H. Morlon, L. K. Nakhleh, B. Oxelman, B. Pfeil, A. Schliep, N. Wahlberg, F. P. Werneck, J. Wiedenhoeft, S. Willows-Munro, and S. V. Edwards. 2019. Embracing heterogeneity: coalescing the Tree of Life and the future of phylogenomics. PeerJ 7.

Browning, S. R. and B. L. Browning. 2011. Haplotype phasing: existing methods and new developments. Nature Reviews Genetics 12:703–714.

Bryant, D., R. Bouckaert, J. Felsenstein, N. A. Rosenberg, and A. RoyChoudhury. 2012. Inferring species trees directly from biallelic genetic markers: Bypassing gene trees in a full coalescent analysis. Molecular Biology and Evolution 29:1917–1932.

Buckler, E. S. and T. P. Holtsford. 1996. *Zea* systematics: Ribosomal ITS evidence. Molecular Biology and Evolution 13:612–622.

Bustamante, E., A. Búrquez, E. Scheinvar, and L. E. Eguiarte. 2016. Population genetic structure of a widespread bat-pollinated columnar cactus. PLOS One 11:e0152329.

Chang, C. C., C. C. Chow, L. Tellier, S. Vattikuti, S. M. Purcell, and J. J. Lee. 2015. Second-generation PLINK: rising to the challenge of larger and richer datasets. GigaScience 4:7.

Chen, J., S. Glemin, and M. Lascoux. 2017. Genetic diversity and the efficacy of purifying selection across plant and animal species. Molecular Biology and Evolution 34:1417–1428.

Chen, J., L. L. Li, P. Milesi, G. Jansson, M. Berlin, B. Karlsson, J. Aleksic, G. G. Vendramin, and M. Lascoux. 2019. Genomic data provide new insights on the demographic history and the extent of recent material transfers in Norway spruce. Evolutionary Applications 12:1539–1551.

Chifman, J. and L. Kubatko. 2014. Quartet inference from SNP data under the coalescent model. Bioinformatics 30:3317–3324.

Copetti, D., A. Búrquez, E. Bustamante, J. L. M. Charboneau, K. L. Childs, L. E. Eguiarte, S. Lee, T. L. Liu, M. M. McMahon, N. K. Whiteman, R. A. Wing, M. F. Wojciechowski, and M. J. Sanderson. 2017. Extensive gene tree discordance and hemiplasy shaped the genomes of North American columnar cacti. Proc. Natl. Acad. Sci., USA 114:12003–12008.

Day, W. H. E. and D. Sankoff. 1987. Computational-complexity of inferring phylogenies from chromosome inversion data. Journal of Theoretical Biology 124:213–218.

De Maio, N., D. Schrempf, and C. Kosiol. 2015. PoMo: An allele frequency-based approach for species tree estimation. Systematic Biology 64:1018–1031.

Degnan, J. H. and N. A. Rosenberg. 2009. Gene tree discordance, phylogenetic inference and the multispecies coalescent. Trends in Ecology and Evolution 24:332–340.

Degnan, J. H. and L. A. Salter. 2005. Gene tree distributions under the coalescent process. Evolution 59:24–37.

Durvasula, A., A. Fulgione, R. M. Gutaker, S. I. Alacakaptan, P. J. Flood, C. Neto, T. Tsuchimatsu, H. A. Burbano, F. X. Pico, C. Alonso-Blanco, and A. M. Hancock. 2017. African genomes illuminate the early history and transition to selfing in *Arabidopsis thaliana*. Proceedings of the National Academy of Sciences of the United States of America 114:5213–5218.

Farris, J. S. 1978. Inferring phylogenetic trees from chromosome inversion data. Systematic Zoology 27:275–284.

Felsenstein, J. 1979. Alternative methods of phylogenetic inference and their interrelationship. Systematic Zoology 28:49–62.

Felsenstein, J. 2004. Inferring Phylogenies. Sinauer Press, Sunderland, MA.

Felsenstein, J. 2005. PHYLIP (Phylogeny Inference Package) version 3.6.

Ferretti, L., E. Raineri, and S. Ramos-Onsins. 2012. Neutrality tests for sequences with missing data. Genetics 191:1397–1401.

Goodman, M., J. Czelusniak, G. W. Moore, A. E. Romeroherrera, and G. Matsuda. 1979. Fitting the gene lineage into its species lineage, a parsimony strategy illustrated by cladograms constructed from globin sequences. Systematic Zoology 28:132–163.

Gramm, J., T. Hartman, T. Nierhoff, R. Sharan, and T. Tantau. 2009. On the complexity of SNP block partitioning under the perfect phylogeny model. Discrete Mathematics 309:5610–5617.

Gusfield, D. 2002. Haplotyping as perfect phylogeny: conceptual framework and efficient solutions. Pages 166–175 in RECOMB ‘02: Proceedings of the Sixth Annual International Conference on Computational biology.

Gusfield, D. 2014. ReCombinatorics: The Algorithmics of Ancestral Recombination Graphs and Explicit Phylogenetic Networks. MIT Press, Cambridge, MA.

Hein, J., M. H. Schierup, and C. Wiuf. 2005. Gene Genealogies, Variation and Evolution: A Primer in Coalescent Theory. Oxford University Press, USA.

Hey, J., Y. J. Chung, A. Sethuraman, J. Lachance, S. Tishkoff, V. C. Sousa, and Y. Wang. 2018. Phylogeny estimation by integration over isolation with migration models. Molecular Biology and Evolution 35:2805–2818.

Hudson, R. R. and N. L. Kaplan. 1985. Statistical properties of the number of recombination events in the history of a sample of DNA-sequences. Genetics 111:147–164.

Huelsenbeck, J. P., J. P. Bollback, and A. M. Levine. 2002. Inferring the root of a phylogenetic tree. Systematic Biology 51:32–43.

Inger, R. F. 1967. Development of a phylogeny of frogs. Evolution 21:369–384.

Junier, T. and E. M. Zdobnov. 2010. The Newick utilities: high-throughput phylogenetic tree processing in the Unix shell. Bioinformatics 26:1669–1670.

Kimura, M. 1969. Number of heterozygous nucleotide sites maintained in a finite population due to steady flux of mutations. Genetics 61:893–903.

Kingman, J. F. C. 1982. On the genealogy of large populations. Journal of Applied Probability 19:27–43.

Korneliussen, T. S., A. Albrechtsen, and R. Nielsen. 2014. ANGSD: Analysis of next generation sequencing data. BMC Bioinformatics 15:356.

Langmead, B. and S. L. Salzberg. 2012. Fast gapped-read alignment with Bowtie 2. Nature Methods 9:357–359.

Lázaro-Nogal, A., S. Matesanz, A. García-Fernández, A. Traveset, and F. Valladares. 2017. Population size, center-periphery, and seed dispersers’ effects on the genetic diversity and population structure of the mediterranean relict shrub *Cneorum tricoccon*. Ecol Evol. 7:7231–7242.

Lee, T. H., H. Guo, X. Y. Wang, C. Kim, and A. H. Paterson. 2014. SNPhylo: a pipeline to construct a phylogenetic tree from huge SNP data. BMC Genomics 15:162.

Li, H., B. Handsaker, A. Wysoker, T. Fennell, J. Ruan, N. Homer, G. Marth, G. Abecasis, R. Durbin, and P. Genome Project Data. 2009. The sequence alignment/map format and SAMtools. Bioinformatics 25:2078–2079.

Liu, L., C. Anderson, D. Pearl, and S. Edwards. 2019. Modern phylogenomics: Building phylogenetic trees using the multispecies coalescent model. Methods in Molecular Biology 1910:211–239.

Ma, B., M. Li, and L. Zhang. 2001. From gene trees to species trees. SIAM J. Comput. 30:729–752.

Maddison, W. P. 1997. Gene trees in species trees. Systematic Biology 46:523–536.

Maddison, W. P. and L. L. Knowles. 2006. Inferring phylogeny despite incomplete lineage sorting. Systematic Biology 55:21–30.

Maddison, W. P. and D. R. Maddison. 2000. MacClade 4: Analysis of phylogeny and character evolution. Sinauer, Sunderland, MA.

Nakhleh, L. 2013. Computational approaches to species phylogeny inference and gene tree reconciliation. Trends in Ecology and Evolution 28:719–728.

Ogilvie, H. A., R. R. Bouckaert, and A. J. Drummond. 2017. StarBEAST2 brings faster species tree inference and accurate estimates of substitution rates. Molecular Biology and Evolution 34:2101–2114.

Page, R. D. M. 1994. Maps between trees and cladistic analysis of historical associations among genes, organisms and areas. Systematic Biology 43:58–77.

Page, R. D. M. and M. A. Charleston. 1997. From gene to organismal phylogeny: Reconciled trees and the gene tree/species tree problem. Molecular Phylogenetics and Evolution 7:231–240.

Pickrell, J. K. and J. K. Pritchard. 2012. Inference of population splits and mixtures from genome-wide allele frequency data. PLOS Genetics 8:e1002967.

Pironon, S., G. Papuga, J. Villellas, A. L. Angert, M. B. García, and J. D. Thompson. 2016. Geographic variation in genetic and demographic performance: new insights from an old biogeographical paradigm. Biological Reviews (Cambridge) 92:1877–1909.

Potts, A. J., T. A. Hedderson, and G. W. Grimm. 2014. Constructing phylogenies in the presence of intra-individual site polymorphisms (2ISPs) with a focus on the nuclear ribosomal cistron. Systematic Biology 63:1–16.

Rannala, B. and Z. Yang. 2017. Efficient Bayesian species tree inference under the multispecies coalescent. Syst. Biol. 66:823–842.

Rheindt, F. E., M. K. Fujita, P. R. Wilton, and S. V. Edwards. 2014. Introgression and phenotypic assimilation in *Zimmerius* flycatchers (Tyrannidae): Population genetic and phylogenetic inferences from genome-wide SNPs. Systematic Biology 63:134–152.

Schmidt-Lebuhn, A. N., N. C. Aitken, and A. Chuah. 2017. Species trees from consensus single nucleotide polymorphism (SNP) data: Testing phylogenetic approaches with simulated and empirical data. Molecular Phylogenetics and Evolution 116:192–201.

Schrempf, D., B. Q. Minh, N. De Maio, A. von Haeseler, and C. Kosiol. 2016. Reversible polymorphism-aware phylogenetic models and their application to tree inference. Journal of Theoretical Biology 407:362–370.

Schrempf, D., B. Q. Minh, A. von Haeseler, and C. Kosiol. 2019. Polymorphism-aware species trees with advanced mutation models, bootstrap, and rate heterogeneity. Molecular Biology and Evolution 36:1294–1301.

Shreve, F. 1951. Vegetation and Flora of the Sonoran Desert vol. 1. Carnegie Institution, Washington, DC.

Springer, M. S. and J. Gatesy. 2016. The gene tree delusion. Molecular Phylogenetics and Evolution 94:1–33.

Sridhar, S., F. Lam, G. E. Blelloch, R. Ravi, and R. Schwartz. 2007. Direct maximum parsimony phylogeny reconstruction from genotype data. BMC Bioinformatics 8:472.

Subramanian, S., U. Ramasamy, and D. Chen. 2019. VCF2PopTree: a client-side software to construct population phylogeny from genome-wide SNPs. PeerJ 7:e8213.

Suh, A., L. Smeds, and H. Ellegren. 2015. The dynamics of incomplete lineage sorting across the ancient adaptive radiation of neoavian birds. PLOS Biology 13:e1002224.

Swofford, D. L. 2002. PAUP*. Phylogenetic Analysis Using Parsimony (*and Other Methods). 4.0 ed. Sinauer, Sunderland, MA.

Swofford, D. L. and S. H. Berlocher. 1987. Inferring evolutionary trees from gene-frequency data under the principle of maximum parsimony. Systematic Zoology 36:293–325.

Than, C. and L. Nakhleh. 2009. Species tree inference by minimizing deep coalescences. PLOS Computational Biology 5:e1000501.

Than, C. and L. Nakhleh. 2010. Inference of parsimonious species trees from multi-locus data by minimizing deep coalescences book section 5, Pages 79–98. Wiley-Blackwell.

Tian, Y. and L. Kubatko. 2017. Rooting phylogenetic trees under the coalescent model using site pattern probabilities. BMC Evolutionary Biology 17:263.

Vachaspati, P. and T. Warnow. 2018. SVDquest: Improving svdquartets species tree estimation using exact optimization within a constrained search space. Molecular Phylogenetics and Evolution 124:122–136.

Wang, J. R. 2013. Analysis and Visualization of Local Phylogenetic Structure within Species. Thesis.

Wang, W. S., R. Mauleon, Z. Q. Hu, D. Chebotarov, S. S. Tai, Z. C. Wu, M. Li, T. Q. Zheng, R. R. Fuentes, F. Zhang, L. Mansueto, D. Copetti, M. Sanciangco, K. C. Palis, J. L. Xu, C. Sun, B. Y. Fu, H. L. Zhang, Y. M. Gao, X. Q. Zhao, F. Shen, X. Cui, H. Yu, Z. C. Li, M. L. Chen, J. Detras, Y. L. Zhou, X. Y. Zhang, Y. Zhao, D. Kudrna, C. C. Wang, R. Li, B. Jia, J. Y. Lu, X. C. He, Z. T. Dong, J. B. Xu, Y. H. Li, M. Wang, J. X. Shi, J. Li, D. B. Zhang, S. Lee, W. S. Hu, A. Poliakov, I. Dubchak, V. J. Ulat, F. N. Borja, J. R. Mendoza, J. Ali, J. Li, Q. Gao, Y. C. Niu, Z. Yue, M. E. B. Naredo, J. Talag, X. Q. Wang, J. J. Li, X. D. Fang, Y. Yin, J. C. Glaszmann, J. W. Zhang, J. Y. Li, R. S. Hamilton, R. A. Wing, J. Ruan, G. Y. Zhang, C. C. Wei, N. Alexandrov, K. L. McNally, Z. K. Li, and H. Leung. 2018. Genomic variation in 3,010 diverse accessions of Asian cultivated rice. Nature 557:43–49.

Xu, B. and Z. H. Yang. 2016. Challenges in species tree estimation under the multispecies coalescent model. Genetics 204:1353–1368.

Xu, D., Y. Jaber, P. Pavlidis, and O. Gokcumen. 2017. VCFtoTree: a user-friendly tool to construct locus-specific alignments and phylogenies from thousands of anthropologically relevant genome sequences. BMC Bioinformatics 18:426.

Yu, Y., T. Warnow, and L. Nakhleh. 2011. Algorithms for MDC-based multi-locus phylogeny inference: Beyond rooted binary gene trees on single alleles. Journal of Computational Biology 18:1543–1559.

Zhang, L. X. 2011. From gene trees to species trees II: Species tree inference by minimizing deep coalescence events. IEEE-ACM Transactions on Computational Biology and Bioinformatics 8:1685–1691.

Zhao, Y. P., G. Y. Fan, P. P. Yin, S. Sun, N. Li, X. N. Hong, G. Hu, H. Zhang, F. M. Zhang, J. D. Han, Y. J. Hao, Q. W. Xu, X. W. Yang, W. J. Xia, W. B. Chen, H. Y. Lin, R. Zhang, J. Chen, X. M. Zheng, S. M. Y. Lee, J. Lee, K. Uehara, J. A. Wang, H. M. Yang, C. X. Fu, X. Liu, X. Xu, and S. Ge. 2019. Resequencing 545 *Ginkgo* genomes across the world reveals the evolutionary history of the living fossil. Nature Communications 10:4201.

Zhu, J. F. and L. Nakhleh. 2018. Inference of species phylogenies from bi-allelic markers using pseudo-likelihood. Bioinformatics 34:376–385.

